# Computational and structural insights into the pre- and post-hydrolysis states of bovine multidrug resistance associated protein 1 (MRP1)

**DOI:** 10.1101/2022.12.27.522006

**Authors:** Ágota Tóth, Veronica Crespi, Angelika Janaszkiewicz, Florent Di Meo

## Abstract

ATP-binding cassette C-family drug membrane transporters play an important role in local pharmacokinetics, *i.e*., drug concentration in cellular compartments. From the structural point of view, only the bovine ortholog of the multidrug-resistance associated protein 1 (*b*MRP1) has been resolved. We here used μs-scaled molecular dynamics simulations to investigate the structure and dynamics of *b*MRP1 in pre- and post-hydrolysis functional states. The present work aims to examine the slight but likely relevant structural differences between pre- and post-hydrolysis states of outward-facing conformations as well as the interactions between MRP1 and the surrounding lipid bilayer. Global conformational dynamics show unfavourable extracellular opening associated with nucleotide binding domain dimerization indicating that the post-hydrolysis state adopts a close-cleft conformation rather than an outward-open conformation. Our present simulations also highlight persistent interactions with annular cholesterol molecules and the expected active role of lipid bilayer in the allosteric communication between distant domains of MRP1 transporter.

## 1. Background

Except for a few members^1–4^, eukaryotic ATP-binding cassette (ABC) superfamily proteins are membrane transporters involved in efflux events, *i.e*., they translocate a myriad of chemically different substrates across the membrane out of cells^1,5^. Among them, the ABCC family plays an active role in pharmacology by transporting xenobiotics such as anionic compounds including glucuro- and sulfo-conjugates^1,6,7^. The so-called multidrug resistance-associated proteins (MRPs) were reported as of importance by the International Transporter Consortium (ITC)^8^ which has stressed^5^ that particular attention must be paid when they are suspected to be major determinants of drug disposition^9^. Indeed, ABCC (dys)functions might be associated with tumour multidrug resistance and drug-drug interactions given the competitive affinity between substrates, with a possible clinical impact.

The structural resolution of ABC transporters has been an active field of research, leading to the elucidation of several conformations, namely inward-facing (IF), outward-facing (OF), and unlock-returned (UR) turnover conformations. These conformations illustrate the alternating access mechanism^10,11^ required for large-scale conformational transitions^1,6,7,12^ along the transport cycle. Recently, structural and functional diversity of ABC transporters has led to a fold-based classification^13^, in which ABCC family is classified as type IV fold^2^. ABCC family members are single proteins comprising two transmembrane domains (TMDs), each consisting of six transmembrane helices (TMHs, Fig. 1). TMDs are connected to evolutionary conserved cytosolic nucleotide binding domains (NBDs). Several bovine ABCC1/MRP1 functional conformations were resolved^6,7,12^ by cryogenic electron microscopy (cryo-EM). Namely, the structures of IF apo (PDB ID: 5UJ9^6^), IF substrate-bound (PDB ID: 5UJA^6^) as well as both the pre- and post-hydrolysis states under OF conformation (PDB IDs: 6BHU^7^ and 6UY0^12^, respectively) were resolved, providing an overview of large-scale transitions occurring along the transport cycle. We recently investigated *b*MRP1 conformations in various lipid bilayer models by molecular dynamics (MD) simulations, suggesting that the NBD asymmetry is key in ATP and substrate-binding events as well as in the allosteric communication between nucleotide binding sites (NBSs) and the substrate-binding pocket^14^. Furthermore, the interplay between *b*MRP1 and lipid bilayer models was also investigated, suggesting an active role of surrounding lipids in the IF-to-OF transition of *b*MRP1, and highlighting direct interactions between phosphatidylethanolamine lipids and cholesterol molecules. Such lipids were shown to actively participate in distance communications between NBSs and the substrate-binding pocket^10,14^. However, focus was mostly paid to the dynamics of milestone conformations along the IF-to-OF transition, prior to ATP-hydrolysis.

**Figure 1.**
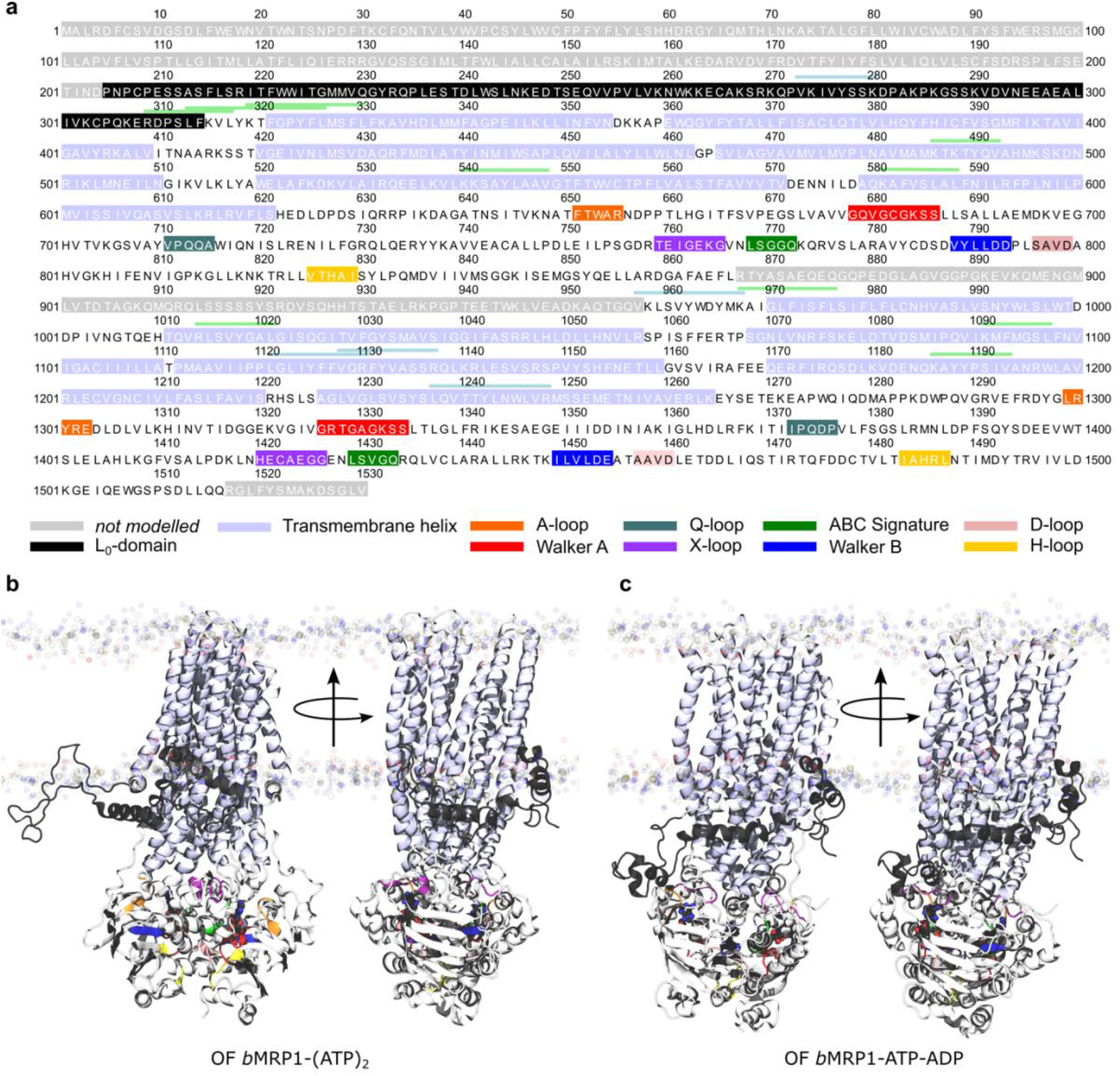
Overview of *b*MRP1 sequence and structures. **a)** Canonical sequence of *b*MRP1 highlighting L_0_ domain and TMHs (respectively in black and light blue). Key NBS motifs are also displayed. Cholesterol Recognition Amino acid Consensus sequence (CRAC) and reverse CRAC (CARC) are depicted respectively in light blue and light green bars. Structures of **b)** OF *b*MRP1-(ATP)_2_ and **c)** OF *b*MRP1-ATP-ADP embedded in a lipid bilayer model using the same colour code. For sake of clarity, CRAC and CARC motifs are not shown here. Only P-, N-atoms of PC and PE polar head groups and O-atom of cholesterol molecules are displayed.

The transport cycle requires the binding of two ATP molecules which are pseudo-symmetrically sandwiched between NBDs. For each NBS, ATP molecule is docked between the A-, H-, D-, and Q-loops, and Walker A and B motifs of one NBD and the ABC signature motif and X-loop of the other NBD^15^ (Fig. 1). MRP1, as other MRPs, is NBS degenerate ABC transporter^15^. The Walker B catalytic glutamate, A-loop aromatic binding tyrosine, and the first ABC signature glycine residues of non-canonical NBS1 are mutated into aspartate (Asp793), tryptophan (Trp653), and valine (Val1431) residues^15^, respectively. This has been proposed to lead to a larger ATP affinity but also to a lower hydrolysis activity, as compared to the consensus NBS2^1,15^. After the substrate release, ATP-hydrolysis has been shown to be key to trigger the OF-to-IF large-scale conformational transition, in which signal transduction from NBDs to TMDs was suggested to occur through coupling helices^7,16^. In NBS degenerate ABC transporters, it is globally accepted that the presence of the functional and canonical NBS2 should ensure the ATP-hydrolysis and subsequent γ-phosphate release suspected to be required to reset the IF conformation^15,17,18^. However, MRP1 cryo-EM resolved structures for pre- and post-hydrolysis functional states do not exhibit significant structural differences. This suggests, altogether with single molecular FRET experiment, that NBD dimer dissociation is the OF-to-IF transition limiting event, but more surprisingly, ATP-hydrolysis and phosphate release might not induce conformational rearrangements in *b*MRP1, in contrast to other ABC transporters^12^. Therefore, this highlights that, even though general trends can be proposed for NBS degenerate ABC transporters, subtle differences between them may lead to different molecular mechanisms^1,15^.

We here use μs-scaled MD simulations to investigate the structure and dynamics of *b*MRP1 in pre- and post-hydrolysis functional states. In line with our previous study, we built an OF *b*MRP1-ATP-ADP model embedded in a simple lipid bilayer model made of POPC (1-palmitoyl-2-oleoyl-sn-glycero-3-phosphocholine), POPE (1-palmitoyl-2-oleoyl-sn-glycero-3-phosphoethanolamine), and cholesterol. The present work aims to investigate the slight but likely relevant structural differences between pre- and post-hydrolysis states of OF conformations as well as the interactions between MRP1 and surrounding lipid molecules. It is important to note that in the present study, since TMD_0_ was not included in our model, TMH labelling follows the short MRP labelling, *i.e*., from TMH1 to TMH12 to ease up comparison with other type IV fold ABC transporters.

## 2. Materials and Methods

### 2.1. Construction of pre- and post-hydrolysis *b*MRP1 models

MD simulations performed for the pre-hydrolysis state, namely OF *b*MRP1-(ATP)_2_, were taken from our previous work^14^. In the present study, the post-hydrolysis OF *b*MRP1-ATP-ADP state was built adopting the exact same protocol^14^. Shortly, initial models were built using the available resolved cryo-EM structures of OF *b*MRP1 bound either to two ATP molecules (OF *b*MRP1-(ATP)_2_, PDB ID: 6BHU^7^) or to ATP and ADP in NBS1 and NBS2, respectively (OF *b*MRP1-ATP-ADP, PDB ID: 6UY0^12^). Engineered EQ mutation experimentally used to lower the rate of ATP-hydrolysis was manually reverted. Since the lack of the so-called TMD_0_ has been shown not to prevent *b*MRP1 transport cycle^6,7,19^, it was not included in the models. The pre-TMH1 lasso domain (L_0_) has been shown to be mandatory for MRP1 function^1,6,20^; however, it was not totally resolved in the cryo-EM structures. Therefore, the missing parts of L_0_ as well as the missing loop between TMH6 and NBD1 were modelled using Modeller v9.23^21^. L_0_ domain was built based on the sequence but also using the IF L_0_ domain as template (Supplementary Table 1). To avoid unphysical steric clashes, final models were shortly minimized in vacuum using Amber package^22,23^ prior to be embedded in the lipid bilayer. Protonation states of titratable residues were assigned by using the PROPKA software^24^, while histidine residues were protonated after careful visual inspection of possible H-bond networks.

OF *b*MRP1 structures were embedded into a lipid bilayer model made of POPC:POPE:Chol (2:1:1), using the CHARMM-GUI input generator^25,26^ and using the *b*MRP1 coordinates from OPM (Orientations of Proteins in Membranes) database^27^ as z-alignment template. The three co-resolved cholesterol molecules were systematically kept in the two structures during all simulations. The original total size of the system was *ca*. 120 x 120 x 180Å^3^ (Supplementary Table 2). To mimic physiological conditions, 0.15 M NaCl salt concentration was used, and the system was solvated with explicit water molecules^28^. The final systems are made of *ca*. 245 000 atoms (Supplementary Table 3).

### 2.2. Molecular dynamics simulations

CHARMM-GUI^25,26^ outputs were converted to Amber format using charmmlipid2amber.py and pdb4amber AmberTools scripts^23,29^. Nucleotides and Mg^2+^ ions were added after building the protein-lipid systems; therefore, neutrality was ensured by randomly removing the corresponding number of counterions. Amber software was used to run simulations, as it was done previously^14^. Protein, lipid, water, and ions were modelled using respectively the FF14SB^30^, Lipid17^31^, TIP3P^28,32,33^ forcefields, and monovalent and divalent ion parameters from Joung and Cheatham^34,35^. ATP and ADP parameters were derived from the Amber database available at http://amber.manchester.ac.uk/^36,37^.

For all simulations, periodic boundary conditions were applied. The cut-off for non-bonded interactions (both Coulomb and van der Waals potentials) was 10 Å. Long-range electrostatic interactions were computed using the particle mesh Ewald method^38^. Amber18 and Amber20 packages^23,29^ using CPU and GPU PMEMD codes were applied for minimization and MD simulations. Each system was initially minimized using a four steps protocol, minimizing: (i) water O-atoms (20000 steps); (ii) all bonds involving H-atoms (20000 steps); (iii) water molecules and counterions (50000 steps) and (iv) the whole system (50000 steps). Thermalization to 310 K was carried out in two steps: (i) under (N,V,T) ensemble conditions, water molecules were thermalized to 100 K during 50 ps using 0.5 fs time integration; (ii) under semi-isotropic (N,P,T) ensemble conditions, the whole system was thermalized from 100 K to 310 K during 500 ps with 2 fs timestep. Each system was finally equilibrated using Berendsen barostat during 5 ns under (N,P,T) ensemble conditions with 2 fs timestep in semi-isotropic conditions. MD production runs were carried out with 2 fs integration timestep under (N,P,T) ensemble conditions with semi-isotropic scaling for 1.5 μs, except for OF *b*MRP1-(ATP)_2_ which reached 2 μs. Langevin dynamics thermostat^39^ with 1.0 ps^-1^ collision frequency was used to maintain the temperature, while either Berendsen barostat^40^ or Monte Carlo barostat^41^ with semi-isotropic pressure scaling were applied to keep the constant pressure set at 1 bar respectively for OF *b*MRP1-(ATP)_2_ and OF *b*MRP1-ATP-ADP systems. The latter was used to speed up computational time after ensuring that system box sizes were sufficiently equilibrated. Snapshots were saved every 100 ps. For each system, three replicas were performed in parallel from the thermalization step to better sample the local conformational space.

### 2.3. Analysis and visualization

The same analysis techniques were used to monitor the simulations as previously^14^. Analyses were performed using the CPPTRAJ^42^ package and in-house Python scripts taking advantage of the MDAnalysis module^43,44^ and the matplotlib v3.3.1 package^45^. Structure visualization and rendering were prepared using the VMD software (v1.9.3 and alpha-v1.9.4)^46^. Analyses were performed only on the equilibrated part of the MD simulations, *i.e*., considering the last 800 or 400 ns shown on root-mean-square deviation (RMSD) profiles reported in Supplementary Fig. 1-2.

The so-called ABC structural parameters^17,47^ were calculated (Supplementary Fig. 3-7) and used to map the ABC conformational space (Fig. 2a) as defined previously^14^, following the work from Hofmann *et al*.^17^ regarding IC angle, EC angle, EC distance and NBD distance as well as the former study carried out by Moradi *et al*. for NBD twist^48^. Shortly, IC angle is defined by the angle between two vectors; both starting from the centre-of-mass of the whole extracellular region and directed toward either the IC region of TMH1, TMH2, TMH3, TMH6, TMH10 and TMH11 or the IC region of TMH4, TMH5, TMH7, TMH8, TMH9 and TMH12. It describes the IC opening of the substrate entry, while the EC angle describes the EC opening for substrate release. EC angle is defined by the angle between two vectors starting from the centre-of-mass of both NBDs and directed toward either the EC region of TMH1, TMH2, TMH9, TMH10, TMH11, and TMH12 or the EC region of TMH3, TMH4, TMH5, TMH6, TMH7, and TMH8. EC distance is defined as the distance between the EC regions of TMH1, TMH2, TMH9, TMH10, TMH11, and TMH12 and the EC region of TMH3, TMH4, TMH5, TMH6, TMH7, and TMH8. NBD distance is defined as the distance between the two NBD centres-of-mass. NBD twist is defined as the dihedral angle between the planes defined by the two NBDs. Residues used to calculate these parameters for resolved ABC transporters are available in supplementary information of Ref.^14^ and values are available in Supplementary Table 4.

**Figure 2.**
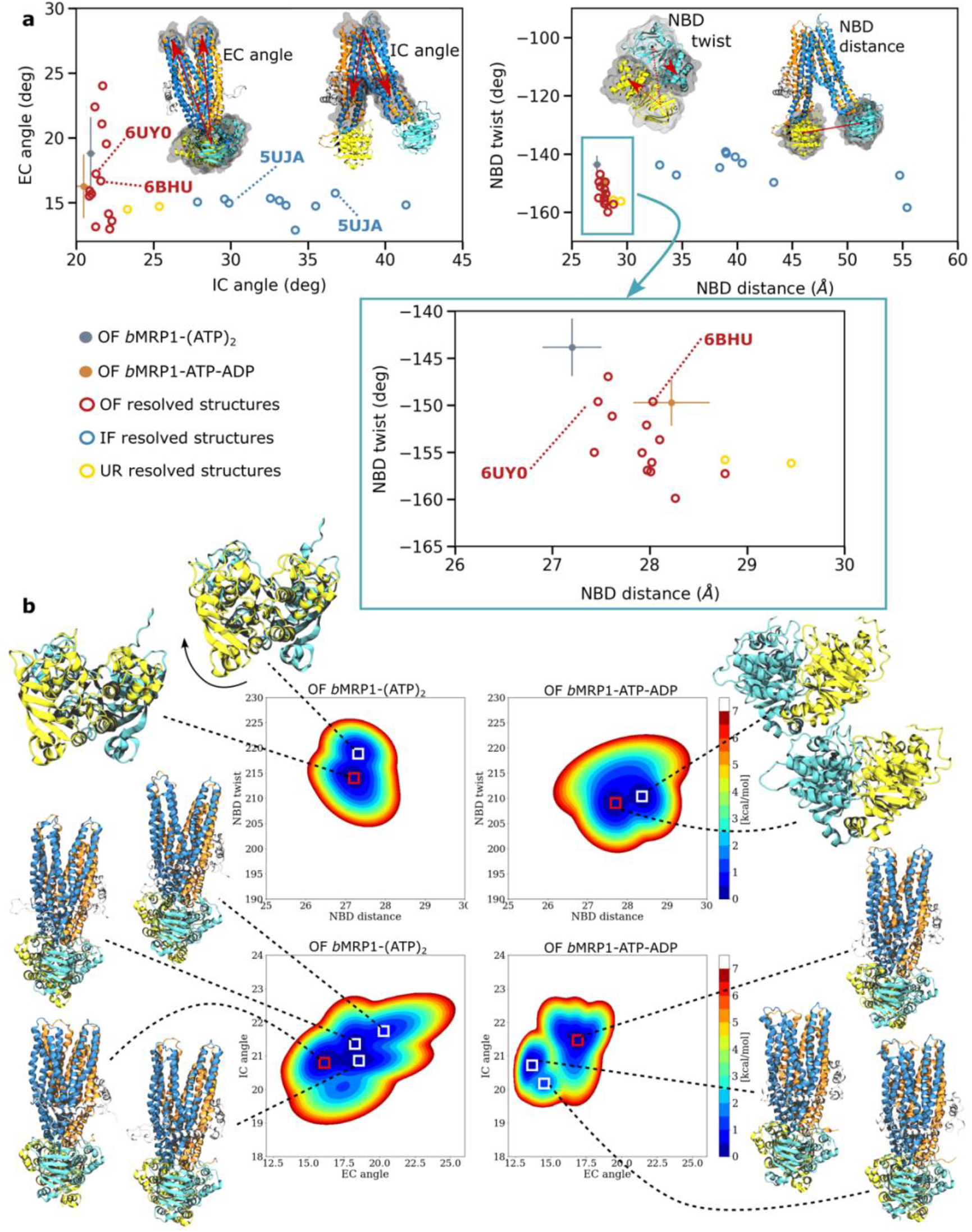
Structural dynamics of OF pre- and post-hydrolysis states. **a)** Projection of *b*MRP1 structural parameters of the pre- and post-hydrolysis states (OF *b*MRP1-(ATP)_2_ and OF *b*MRP1-ATP-ADP, respectively) onto the ABC conformational space obtained from resolved ABC structures. The last 800ns from each replica were considered and averaged. PDB IDs of the resolved *b*MRP1 cryo-EM structures are explicitly mentioned. The zoom shows the OF area of the NBD twist – NBD distance diagram. The first and second TMDs are respectively depicted in orange and blue, and NBD1 and NBD2 are respectively coloured yellow and cyan. **b)** System-dependent local conformational landscapes obtained from the GMM-based approach developed in the InfleCS method (See Refs. ^49,50^) highlighting the influence of nucleotides on *b*MRP1 structural dynamics. Representative snapshots for each basin are shown. The most populated basins are depicted by red square on the free energy surfaces.

The GMM-based InfleCS^49,50^ clustering method was used to map the subspace conformation free energy surfaces (Fig. 2b). The grid size was set at 80 point per variable, and 2 to 12 gaussian components for each GMM were considered to fit densities. GMM functions were obtained by considering a maximum of 20 iterations. Free energy surfaces were calculated using the ABC structural parameters as structural determinants. Since the relevance of InfleCS^49,50^ approach strongly relies on the quality of sampling during MD simulations, InfleCS^49,50^ here only pictures the free energy landscape around the sampled local minima. As previously achieved^14^, the relevance of InfleCS for OF *b*MRP1-ATP-ADP was monitored by calculating convergence profiles for each structural parameter separately (Supplementary Fig. 8). H-bond analyses (Fig. 4c) using 3.5 Å and 120° cut-offs respectively for distance and angle were performed by CPPTRAJ^42^. Contacts between the nucleotides and their binding site were calculated (Supplementary Fig. 9). Leaflet dependent lipid densities were calculated (Supplementary Fig. 10-11) as previously^14^, *i.e*., using the grid keyword in CPPTRAJ^42^ concentrating on polar head lipids (PE or PC) or cholesterol OH-group. Focus was paid only to the high-density polar head region obtained from lipid bilayer models^46,51^.

**Figure3.**
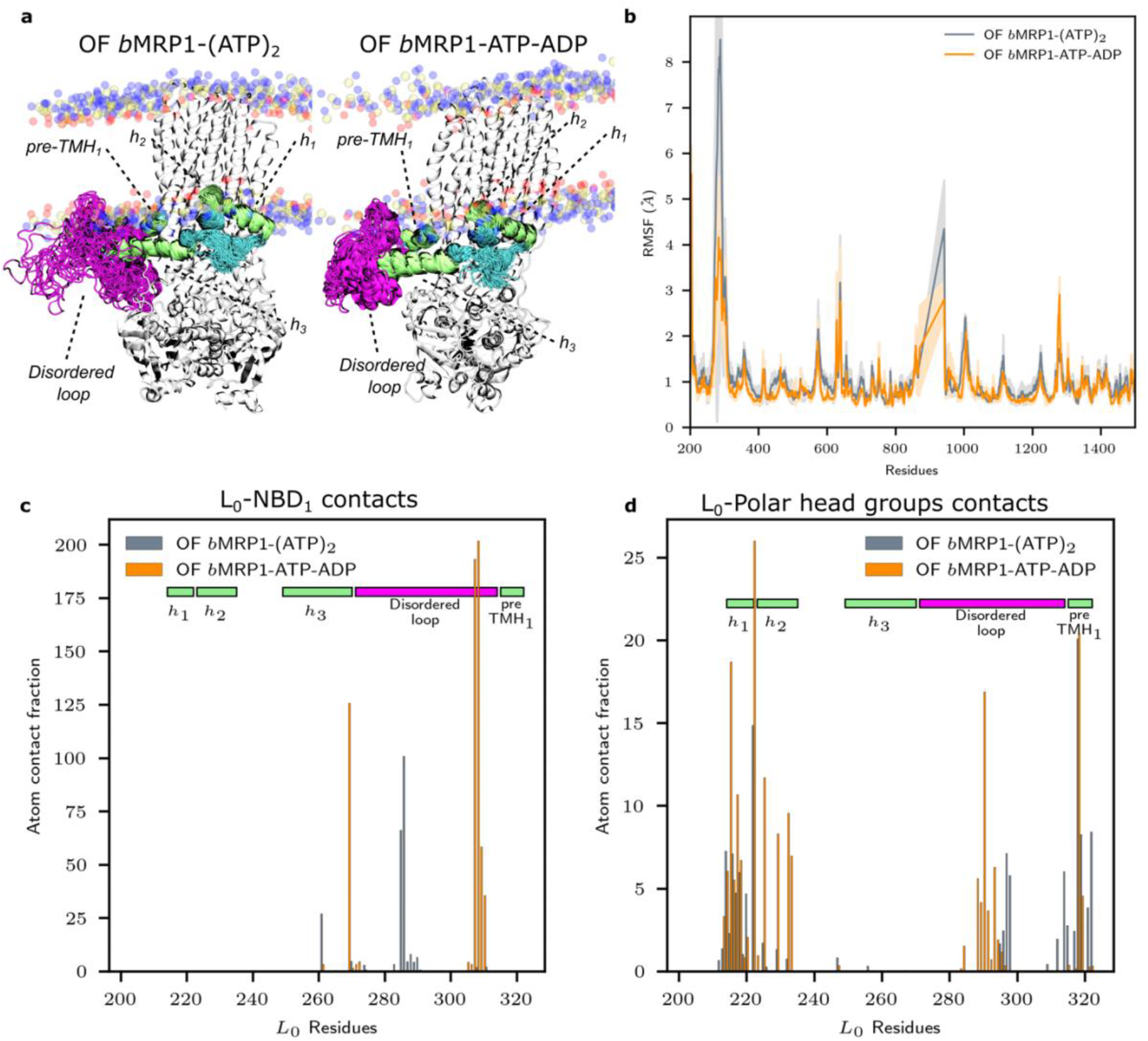
Putative dynamics and interactions of L0 in ^ΔTMD0^MRP1 model. **a)** Representation of the OF *b*MRP1 pre- and post-hydrolysis structure highlights the L_0_-dynamics. 75 L_0_-domain structures taken along the MD simulations are displayed. L_0_ α-helices and long disordered loops are depicted green and purple, respectively. **b)** Average per-residue RMSF calculated over the last 400 ns of each replica. Atomic contact fractions over the last 400ns between the L_0_ domain and **c)** NBD1 and **d)** lipid polar heads groups and cholesterol molecules.

**Figure 4.**
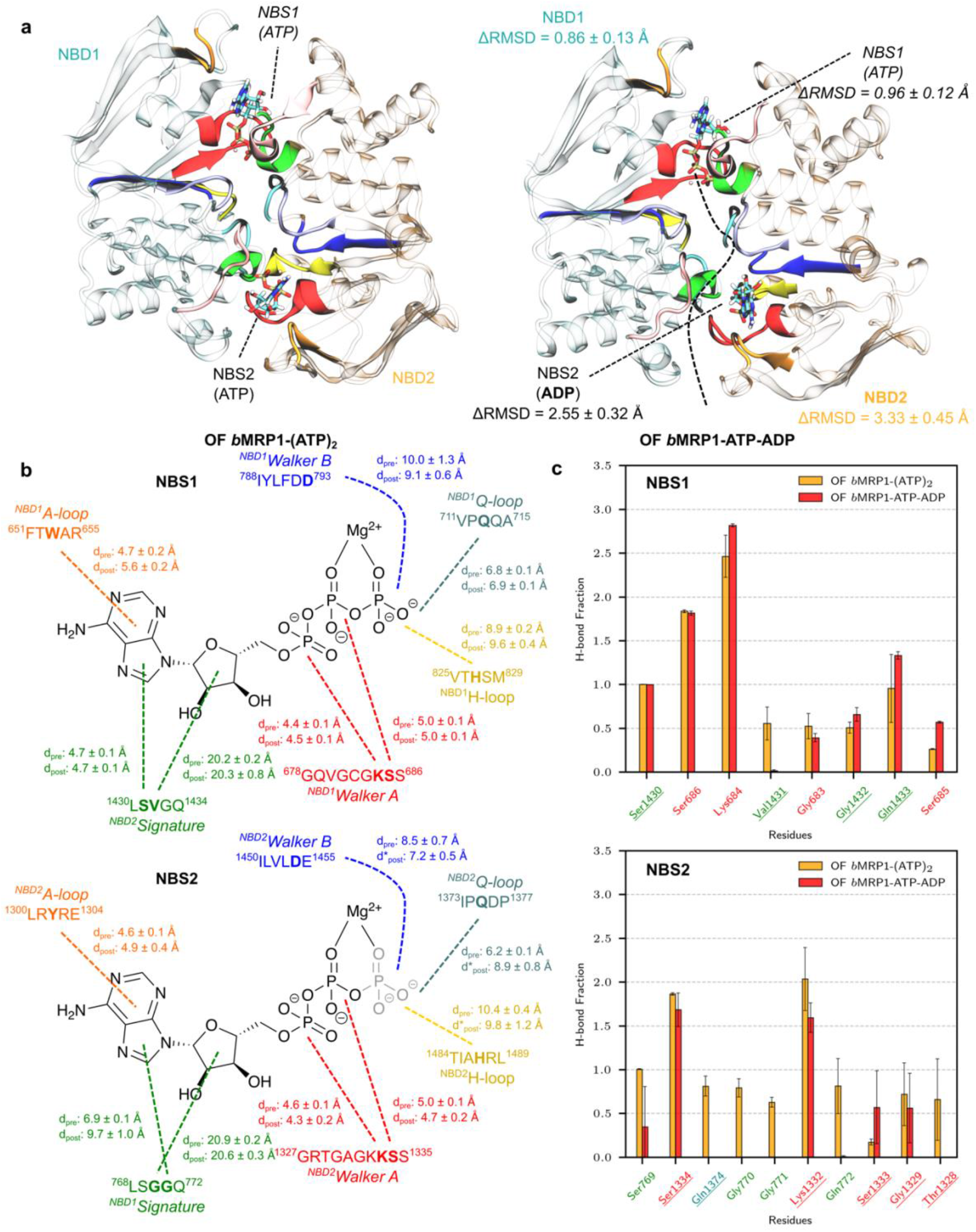
Nucleotide-binding sites in the pre- and post-hydrolysis states of OF *b*MRP1. **a)** ΔRMSD of NBD1, NBD2 and NBS1, NBS2 of the post-hydrolysis state compared to the pre-hydrolysis state. The post-hydrolysis state is slightly more open as it is indicated by the dashed line. Conserved motives are coloured as follows: Walker A red, Walker B blue, signature motif green, A-loop orange, Q-loop teal, X-loop pink, D-loop cyan, and H-loop yellow. **b)** Important distances between the nucleotides and key residues in the nucleotide-binding sites. **c)** Calculated H-bond networks between nucleotides and NBS1 (top) or NBS2 (bottom) in OF *b*MRP1-(ATP)_2_ and OF *b*MRP1-ATP-ADP.

Taking inspiration from previous work performed on the ABCB1/P-gp transporter^52^, *b*MRP1 sequence was scanned for Cholesterol Recognition Amino acid Consensus (CRAC) motifs as well as reverse CRAC (CARC) motifs, using the ScanProSite tool^53^, available in the Swiss Bioinformatics Resource Portal Expasy. CRAC and CARC motifs were respectively defined as (Leu/Val)-X_1-5_-(Tyr)-X_1-5_-(Lys/Arg) and (Lys/Arg)-X_1-5_-(Phe/Tyr)-X_1-5_-(Leu/Val) sequences. X refers to as any amino acid. CRAC motif was extended to its initial central tyrosine definition to sample more sequences.

Allostery network pathways, as previously^14^, were determined using the Allopath approach developed by Westerlund *et al*.^54,55^; which evaluates the distant communication between different domains by monitoring the information flow along complex molecular network for which each protein residue is considered as a node. The so-called interactors (*i.e*., nucleotides, lipids, and magnesium) can also be considered in the network. For each lipid, three nodes were assigned for polar head group, sn_1_- and sn_2_-tails. Likewise, nucleotides were split into three nodes, namely adenyl and ribosyl moieties and phosphate tail. Magnesium atoms and cholesterol molecules were considered as single nodes. Allosteric communications were calculated between two distant domains by calculating a node contact map as well as the mutual information matrix. The latter models the correlation between nodes considering protein residue and interactor nodes. While the network interaction analysis approach^56^ focuses on deciphering the optimized allosteric pathway between two nodes, the Allopath tool^54,55^ extends this idea by accounting for all possible pathways at once including surrounding interactors such as lipids. The overall efficiency of information flow between sinks and sources can then be estimated from the current flow closeness while the “involvement” of each protein residue in the distance communication can be evaluated from the current flow betweenness, providing hints about the plausible different pathways. Allosteric communications were calculated from extracellular regions (TMH1, 2, 10, and 11 and TMH4, 5, 7, and 8) to each NBS A-loop aromatic residues as defined in Supplementary Table 5.

## 3. Results

### 3.1. MD simulations reveal slight but relevant structural differences between the pre- and post-hydrolysis states

Conformational spaces sampled for pre- and post-hydrolysis states (*i.e*., OF *b*MRP1-(ATP)_2_ and OF *b*MRP1-ATP-ADP) were examined focusing on key ABC structural parameters (Fig. 2a). NBD distances and intracellular (IC) angles were used to monitor the intracellular opening, while the extracellular angle (EC) was used to assess the opening towards the external cell compartment. NBD rocking-twist as originally defined by Moradi *et al.^47^* was also monitored. These parameters were projected onto the ABC conformational space defined by measuring ABC structural parameters of a large data set of experimentally resolved ABC transporters (Fig. 2a, and Supplementary Table 4). The so-called IF, OF, and asymmetric UR turnover^17^ states are explicitly separated using these structural parameters. For both pre- and post-catalytic states, IC angle shows similar trends picturing small IC opening, values ranging from 19.4 to 22.6°. Furthermore, they did not show large variability suggesting stable conformations (Fig. 2a, Supplementary Fig. 3). On the other side of the membrane, post-hydrolysis state exhibits smaller EC angle values than pre-catalytic state, ranging from 13.8 to 23.6 and from 12.6° to 19.3°, respectively for pre- and post-catalytic states. This is in line with the expected sealing of EC gate which is favoured after ATP-hydrolysis. EC opening variability is larger than for IC angle depicting the expected NBD locked conformation (Fig. 2a, Supplementary Fig. 4-5).

Pre- and post-catalytic states mostly differ in the NBD dimer arrangement. The NBD rocking-twist angle of the post-hydrolysis state does not deviate from the originally resolved structure, while the NBD distance value is slightly larger (Fig. 2a, Supplementary Fig. 6-7). The opposite trend was observed in pre-catalytic state for which NBDs get closer to each other in MD simulations with respect to the cryo-EM resolved structure. This might be explained by stronger NBD-NBD non-covalent interactions in biomimetic membranes compared to detergent, as suggested in our previous work^14^.

Furthermore, the E1454Q mutation used for cryo-EM resolution may also alter ATP-binding in NBS2. Interestingly, the post-catalytic OF *b*MRP1-ATP-ADP state exhibits a lower NBD twist than the pre-catalytic state by *ca*. 5°. Noteworthy, the post-hydrolysis state which is expected to be before the OF-to-IF transition is getting closer to the conformational space of the UR structures which were resolved in native nanodisk environment. This is in agreement with the recent structural findings suggesting that ABC transporter might adopt UR state prior to get back to IF conformational state, resetting the transport cycle^17^. However, the use of NBD distances should be carefully considered, especially, for NBD degenerate ABC transporters such as MRP1, given the expected pivotal role of ATP-bound NBS1 in maintaining NBD dimerization along the transport cycle^15^, in agreement with our previous findings^14^.

Assuming that our MD simulations were long enough to independently sample the local conformational subspace of pre- and post-catalytic metastable states, local free energy surfaces were calculated according to ABC structural parameters using the InfleCS framework^49,50^ (Fig. 2b, Supplementary Fig. 8). Representative snapshots considering the different basins are reported in Fig. 2b. Interestingly, two main populations were captured from MD simulations performed on the post-catalytic OF *b*MRP1-ATP-ADP state regarding NBD distance, showing a slightly larger flexibility in terms of NBD opening while it is unlikely to happen in the pre-catalytic state. Likewise, a small difference for NBD rocking-twist parameter between pre- and post-hydrolysis states is captured from the free energy point of view, confirming that ATP-hydrolysis shifts the free energy surface toward more open and less twisted NBD dimer conformation.

On the other side of the membrane, pre-hydrolysis state simulations show more flexible EC opening compared to the post-catalytic state, as shown by the four different representative snapshots (Fig. 2b). However, the most likely population remains the most EC closed conformation. MD simulations performed on OF *b*MRP1-ATP-ADP state surprisingly propose two well-separated populations. Interestingly, the transition from one to the other requires crossing a relatively high energy barrier (*ca*. 3 kcal.mol^-1^). Our simulations suggest that such transition may be correlated with slightly more closed IC gate. However, this should be carefully considered and confirmed by further investigations. Indeed, the two populations exhibiting smaller EC opening are not the most likely and were obtained from a single replica. Therefore, no exchange between the two basins were observed during our MD simulations.

### 3.2. Structural investigations of L_0_-domain in ^ΔTMD0^MRP1 models

Particular attention was also paid to the lasso domain (L_0_) which connects the TMD_0_ and the functional core of MRP1. TMD_0_ was not included in the present MRP1 models since it has been shown that its absence does not affect the transport function^6,7,19^.

However, experiments stressed out that L_0_-less MRP1 is not functional^1,6,20^. Even though the structure of L_0_ remains unclear, cryo-EM structures as well as MD simulations suggest that L_0_ consists in three short α-helices in the N-terminal region (namely h1, h2 and h3, from Asn203 to Lys267) followed by a large flexible disordered loop (Ser268-Lys307) connected to the pre-TMH1 elbow helix (Fig. 3a). Globally, MD simulations reveal large flexibility of the L_0_ domain (Fig. 3a-b) as pictured by comparing RMSD for the models along the simulations with and without considering L_0_ region (Supplementary Fig. 1-2). Indeed, assessing RMSD only on the ABC core (*i.e*., excluding L_0_) showed much smaller deviations along the simulations of pre- and post-catalytic states (Supplementary Fig. 2). Domain flexibilities were also assessed by measuring the per-residue root-mean-square fluctuations (RMSF) (Fig. 3b). Interestingly, h1, h2 and h3 helices remain stable along MD simulations, showing stable dynamics while L_0_ disordered domain exhibits higher flexibility (Fig. 3a-b). The Asn203-Lys267 region does not exhibit large flexibility, remaining in close contact with pre-TMH1, TMH1, TMH2, TMH10, and TMH11. Interestingly, in both pre- and post-catalytic states, L_0_ flexibility suggests two distinct behaviours regarding the Ser268-Lys307 disordered domain (Fig. 3a). In the post-hydrolysis state, the disordered loop appeared less flexible than in the pre-hydrolysis state. We here propose that the lower flexibility may be due to weak interactions with NBD1 as shown by contact analyses (distance < 8.0 Å) in Fig. 3c-d. Even though L_0_ roughly maps the same subspace in the pre- and post-hydrolysis states, more contacts were observed between L_0_ disordered domain and NBD1 in the post-hydrolysis state. Our results suggest that L_0_ conformation may differ after ATP hydrolysis, leading to local rearrangements of L_0_. Likewise, more important contacts between L_0_ and lipid polar head groups were also observed in the post-hydrolysis state (Fig. 3d). However, these results should be considered carefully since: (i) TMD_0_ was not modelled while it might reduce the overall flexibility of L_0_ given its expected membrane anchoring role; (ii) L_0_ is expected to undergo phosphorylation by post-translational modifications which may strongly affect non-covalent interactions with its environment (*i.e*., NBD1 and lipid polar heads).

### 3.3. Differences in the nucleotide-binding of the pre- and post-hydrolysis states

As expected, given the MD simulation timescale precludes the investigation of ADP release, no NBD dimer dissociation event was observed in any of the present MD simulations for the post-catalytic state (Fig. 2a). As NBD distance and NBD twist values showed differences in the pre- and post-hydrolysis states, special attention was paid to the NBSs. MRP1 belongs to NBD degenerate ABC transporters of which NBS1 is expected to exhibit (i) increased important ATP-binding but (ii) significantly lower ATP-hydrolysis rate. Nucleotide-bound state differences between pre- and post-hydrolysis states are expected to modulate the NBD dimer supramolecular arrangement. In this context, NBD2 arrangement with respect to NBD1 was evaluated by calculating ΔRMSDs focusing on NBDs either separately or using NBD1 position as a reference over the trajectories. Interestingly, NBD1 secondary structure is conserved during the hydrolysis (ΔRMSD being 0.86 ± 0.13 Å, Fig. 4a). NBD2 secondary structure deviates slightly more (ΔRMSD being 3.33 ± 0.45 Å, Fig. 4a) while comparing pre- and post-hydrolysis states. However, NBD1-aligned structure clearly highlights the structural deviation of post-catalytic NBD dimer as compared to pre-catalytic state. The pivotal role of NBS1 is confirmed as well as the asymmetric NBD motion along the transport cycle^15^. For instance, NBS2 exhibits larger ΔRMSD value, (2.55 ± 0.32 Å) as compared to NBS1 (0.96 ± 0.12 Å).

Moreover, visual inspection clearly underlines the opening of NBD dimer on the side of NBS2 in line with ABC parameter projections (Fig. 2a), as previously suggested^14^.

Local key distances proposed to maintain the nucleotide in the binding site^14,57^ were also monitored between either ATP or ADP molecules and well-known conserved motifs (Fig. 4b). Distances between ATP molecules and NBS1 binding motifs are globally conserved when comparing pre- and post-hydrolysis states. This is in agreement with former observations made on NBD degenerate ABC transporters suggesting that ATP remains constantly bound in NBS1 along several transport cycles^15^. In contrast, distances between the nucleotide and NBS2 motifs exhibit larger variabilities between pre- and post-hydrolysis states, especially between adenosyl and ABC signature motif and between diphosphate moiety and Q-loop, H-loop, or Walker B motifs. The most important deviation was observed with the Q-loop motif (respectively 6.2 ± 0.1 and 8.9 ± 0.8 Å in pre- and post-hydrolysis states). This might be easily explained by the absence of γ-phosphate which is supposed to bind the Q-loop motif. However, in presence of ADP in NBS2, motifs supposed to bind γ-phosphate tend anyway to locally reshape by getting closer to β-phosphate moiety (*e.g*., ABC signature, H-loop, and Walker B), as shown by distances measured with terminal β-phosphate O-atom (d*_post_ in Fig. 4b). This suggests a local rearrangement of NBDs after ATP-hydrolysis.

More specifically, binding modes of the nucleotides were also investigated by assessing atomic contacts, H-bond networks as well as van der Waals and Coulomb potentials between ATP or ADP molecules and NBS motifs (Fig. 4cb, Supplementary Fig. 9, 12). It must be stressed that van der Waals and Coulomb potentials can only provide qualitative hints to compare the binding sites due to a compensation of errors between the two potentials especially at short distances^58^. H-bond fractions (Fig .4c) highlight the significant difference between pre- and post-hydrolysis H-bond network in NBS2, especially regarding the ABC signature motif and Q-loop. This is confirmed by interatomic contact analyses since there are significantly less contacts in the post-hydrolysis state in NBS2 (Supplementary Fig. 9). Walker A motif in NBS2 has slightly looser connections with ADP than ATP, which might be involved in the subsequent nucleotide release. NBS-nucleotide atomic contact fraction maps confirm weaker interactions between ADP and NBS2 ABC signature motif and Q-loop, while contacts remain similar with other motifs (Supplementary Fig. 9). Interestingly, ABC signature motif has similar H-bond fraction in NBS1 in pre- and post-hydrolysis states. NBS1 Walker A motif, that binds α- and β-phosphates, exhibits slightly stronger connections in the post-hydrolysis state in line with the expected docking of ATP molecule along multiple transport cycles^15^.

Mg^2+^-binding modes were also investigated in NBS1 and NBS2 regarding pre- and post-hydrolysis states for which significant differences were observed after γ-phosphate release (Fig. 5 and Table 1). In agreement with literature and cryo-EM resolved ATP-bound structures^7,12,57^, key Mg^2+^-binding residues were observed for ATP-binding in NBS, *i.e*., Q-loop glutamine, Walker A serine as well as ATP phosphate groups (Fig. 5). These key distances remain stable along MD simulations (Table 1). Similar trends were observed for the ATP-binding in NBS1 of OF *b*MRP1-ATP-ADP. Only the distance between α-phosphate and Mg^2+^ exhibited slightly larger value (3.11 ± 0.05 and 4.36 ± 0.09 in OF *b*MRP1-(ATP)_2_ and *b*MRP1-ATP-ADP, respectively). Interestingly, our simulations suggested that the γ-phosphate release locally reshapes Mg^2+^-binding in NBS2. Surprisingly, in contrast to resolved cryo-EM structure^12^, MD simulations showed the participation of Walker B anionic residues in Mg^2+^-binding (Asp1453 and catalytic Glu1454) contrary to ATP-binding (Fig. 5). However, Q-loop glutamine and Walker A serine residues remained in close vicinity to Mg^2+^ participating in its binding but to a less extent than in ATP-binding as proposed by the higher variabilities calculated along MD simulations (Table 1).

**Figure 5.**
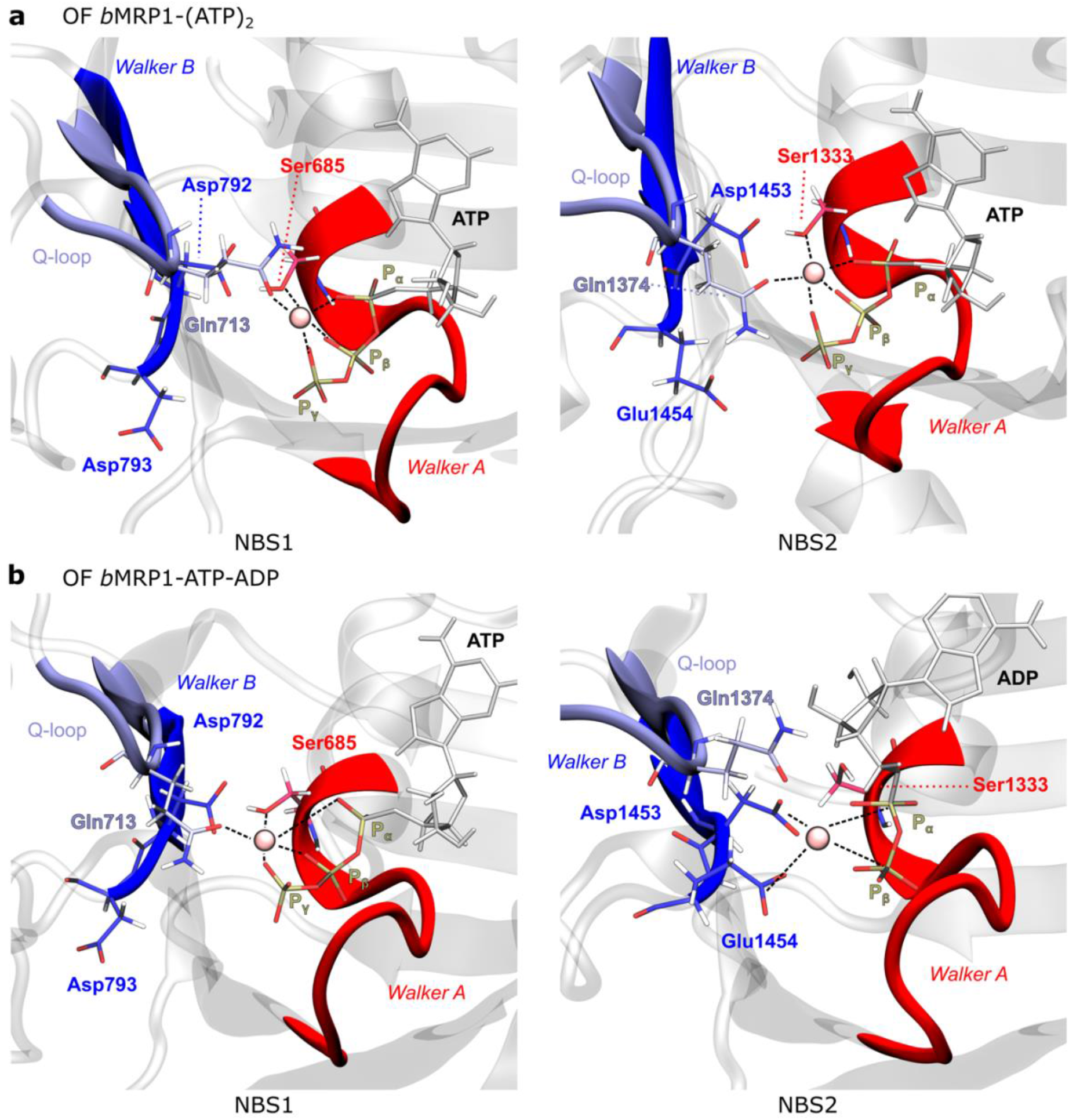
Binding modes of the Mg^2+^ ions. Representative snapshots highlight the binding of magnesium cation by nucleotide phosphate, Walker B (blue), Q-loop (teal) and Walker A (red) from MD simulations performed on NBS1 and NBS2 **a)** OF *b*MRP1-(ATP)_2_ and **b)** OF *b*MRP1-ATP-ADP. Binding distances are shown in black dashed lines. Key residues are also reported. Average values are reported in Table 1.

**Table 1.** Average key distances (Å) between Mg^2+^ atom and surrounding binding residues from MD simulations.

| Motif/Residue |  | NBS1 |  | NBS2 |  |
| --- | --- | --- | --- | --- | --- |
|  |  | *bMRP1*-(ATP)<sub>2</sub> | *bMRP1*-ATP-ADP | *bMRP1*-(ATP)<sub>2</sub> | *bMRP1*-ATP-ADP |
| ATP | Pα | $3.11 \pm 0.05$ | $4.36 \pm 0.09$ | $3.15 \pm 0.05$ | $3.26 \pm 0.40$ |
| | Pβ | $2.99 \pm 0.05$ | $3.15 \pm 0.05$ | $3.01 \pm 0.05$ | $2.86 \pm 0.26$ |
| | Pγ | $3.02 \pm 0.06$ | $3.13 \pm 0.06$ | $3.10 \pm 0.06$ | - |
| Walker B | Asp792/Asp1453 | $5.79 \pm 1.74$ | $4.25 \pm 0.21$ | $5.31 \pm 1.03$ | $3.22 \pm 1.27$ |
| Walker B | Asp793/Glu1454 | $10.29 \pm 1.67$ | $8.44 \pm 0.96$ | $7.73 \pm 0.94$ | $2.92 \pm 0.10$ |
| Q-loop | Gln713/Gln1374 | $2.02 \pm 0.04$ | $2.03 \pm 0.06$ | $2.05 \pm 0.04$ | $3.43 \pm 1.31$ |
| Walker A | Ser685/Ser1333 | $2.04 \pm 0.04$ | $2.11 \pm 0.07$ | $2.06 \pm 0.04$ | $3.74 \pm 1.12$ |

### 3.4. Protein-Lipid interactions

We have recently shown that lipid bilayer composition only slightly affects the IF state structures close to the local equilibrium subspace^14^. By considering here only one lipid bilayer model, we focused on the interplay between surrounding lipids and *b*MRP1 OF structures. Lipid distribution around *b*MRP1 structures was calculated by considering all trajectories. Lower and upper leaflet 2D densities of PE groups and cholesterol were evaluated (Supplementary Fig. 10-11). We also identified regions in which PE and cholesterol may likely preferentially bind to *b*MRP1 TMHs (Fig. 6a).

**Figure 6.**
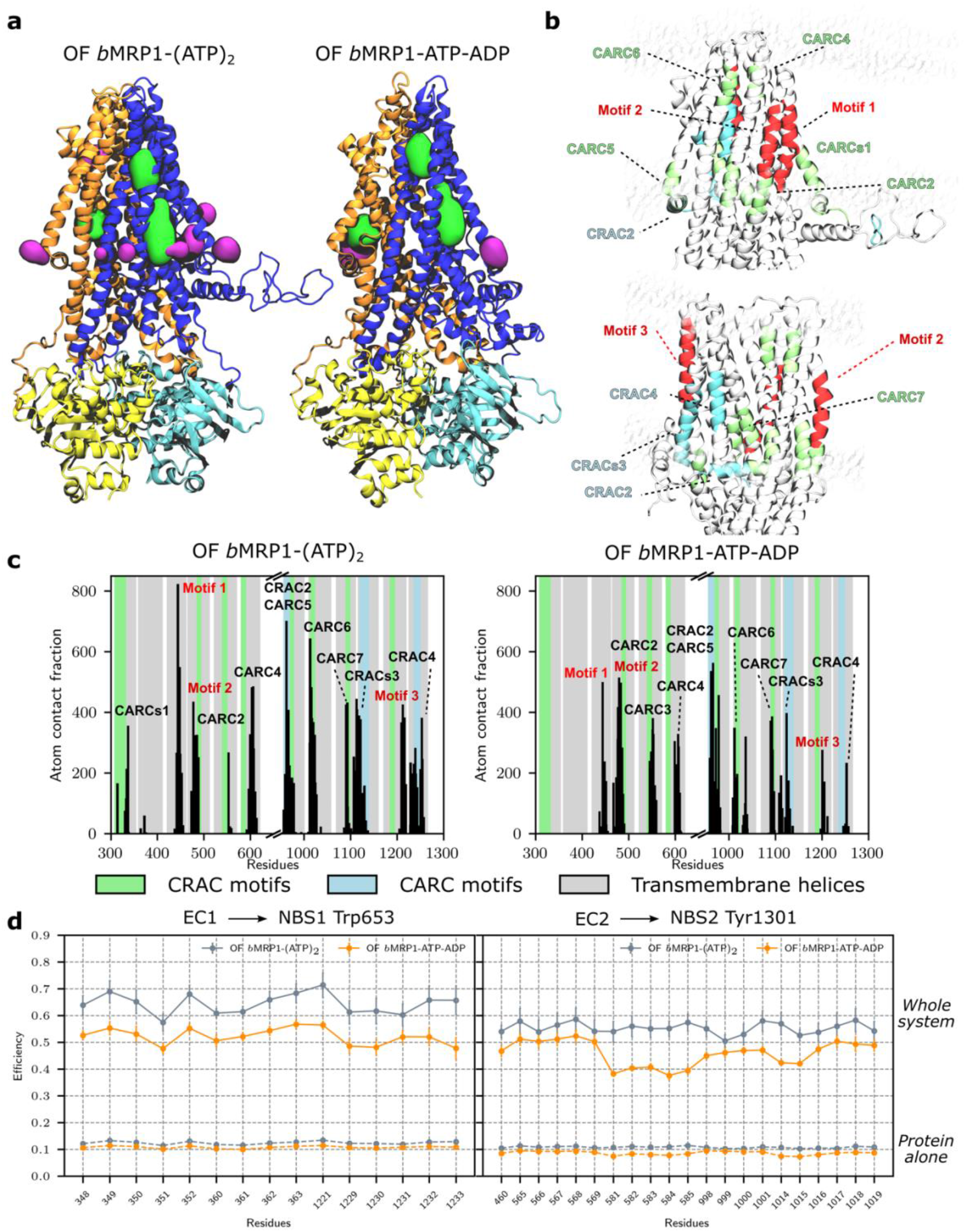
The role of the lipids on the dynamics of the pre- and post-hydrolysis states. **a)** Key cholesterol (green), and PE lipid (magenta) hotspots defined by presence likelihood higher than 50%. **b)** Location of identified CARC (green) and CRAC (blue) motifs from sequence scanning as well as motifs defined from MD simulations. **c)** Atomic contact map between cholesterol and *b*MRP1 residues highlighting preferential binding regions. Predicted CARC and CRAC motif as well TMH are coloured in green, blue and grey. **d)** Allosteric efficiency of the information flow from the EC regions to the NBS A-loop aromatic residues.

Plausible PE binding site shed light on key PE-protein H-bond interactions and thus hotspots that are mainly observed in the lower leaflet. This is in agreement with the known *in vivo* asymmetry for cell membrane composition, since PE lipids are mostly located in the lower leaflet than the upper one. PE distributions suggest that PE-binding is more likely prior to ATP-hydrolysis since more hotspots were observed in OF *b*MRP1-(ATP)_2_ than in OF *b*MRP1-ATP-ADP simulations. However, such assumptions should be considered carefully given that the present timescale is not sufficient to provide an accurate sampling of surrounding lipid motions. Cryo-EM OF structures were resolved including three cholesterol molecules that were suggested to actively participate in the allosteric communication between substrate-binding pocket and NBD^14^.

As observed for the pre-hydrolysis state, resolved cholesterol remains in close contact with pre-TMH7 elbow helix all along the simulation (Fig. 6a). To document about cholesterol binding events, we scanned the *b*MRP1 sequence to identify cholesterol recognition amino acid consensus (CRAC) motifs and reverse CRAC (CARC) motifs. By excluding irrelevant CRAC and CARC motifs located out of the *b*MRP1 TMDs, 8 CRAC and 4 CARC motifs were identified and located in different *b*MRP1 domains (Table 2 and Fig. 6b). The three first CARC motifs (namely CARC1a-c) were assigned to a single CARCs motif since (i) they are overlapping in the *b*MRP1 sequence (Fig. 1a) and (ii) are located in the pre-TMH1 region that has been shown to be a preferential cholesterol binding site in *b*MRP1 IF states^14^. Likewise, CRAC3a and CRAC3b overlap leading to a potential larger binding motif from Leu1121 to Arg1137 in TMH10. To confirm their relevance with potential cholesterol binding, close contact analyses (distance < 4.5 Å) between cholesterol molecules and *b*MRP1 were performed (Fig. 6c) since it has been shown that the use of CRAC/CARC motifs is not sufficient to extensively sample cholesterol recognition domains^52^. Most of the identified CRAC and CARC motifs exhibit large contacts confirming their role in cholesterol-protein interactions. Only CARC8 and CRAC1/4 did not show significant contacts during MD simulations.

**Table 2.**
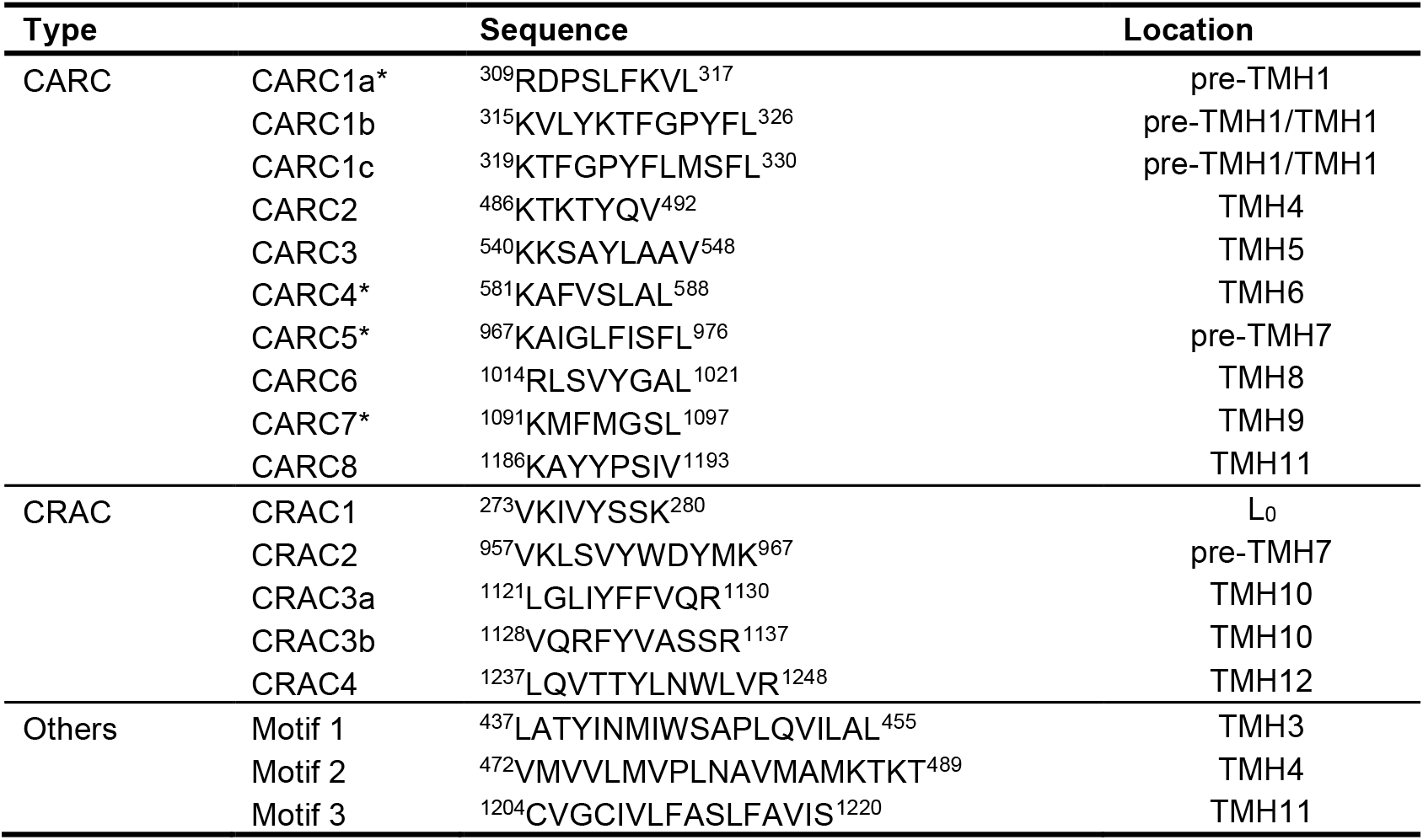
Location of identified CRAC and CARC motifs within the transmembrane region of *b*MRP1 as well as other amino acid sequences associated with high contacts from present MD simulations (namely Motifs 1, 2 and 3). CRAC-like motifs (*i.e*., with central phenylalanine) are highlighted with *.

CRAC1 is located in L_0_ at the interface with lipid bilayer while CARC8 is located in TMH11. MD simulations also suggested that three other motifs may play a role in binding cholesterol molecules (Table 1 and Fig. 6c), namely Motif 1, 2 and 3 which are located in TMH3, 4 and TMH11. Motifs 2 and 3 are adjacent respectively to CARC2 and CARC8. Interestingly, Motif 1 sequence exhibits an incomplete CRAC motif, missing only the ending cationic residue (^437^LATYINMIW^445^). Such incomplete pattern was also suggested for P-gp^52^. Motif 1 and motif 2 are involved in one persistent resolved cholesterol binding along MD simulations (Fig. 6a). MD simulations suggest that most of the cholesterol contacts are conserved during the ATP-hydrolysis event given that similar contact patterns were obtained for OF *b*MRP1-(ATP)_2_ and *b*MRP1-ATP-ADP.Only the cholesterol binding event at the pre-TMH1/TMH1 region (CARCs1) was not observed in the post-hydrolysis *b*MRP1-ATP-ADP state.

Given that annular lipids were shown to play an active role in distance communication for membrane proteins including P-gp^59^ and *b*MRP1^14^, the efficiency of the information flow between EC regions to NBS A-loop aromatic residues (namely Trp653 and Tyr1301 for NBS1 and NBS2, respectively) was evaluated by calculating the so-called current flow closeness using the Allopath tool^54,55^. Distant communications were monitored by considering the two distinct EC regions from TMH bundles (*i.e*., EC1 and EC2 regions respectively for extracellular regions of TMH1, 2, 10, and 11 and TMH4, 5, 7, and 8) since they were shown to be responsible for the EC opening^**17**^. As previously reported^10,14^, MD simulations confirmed the central role of membrane in participating in the allosteric communication between cytosolic domains and EC region of TMHs (Fig. 6d, Supplementary Fig. 13-14). The presence of surrounding lipids significantly increased information flow efficiencies confirming that surrounding lipids actively participate in the allosteric pathway along the transporter. This was also pictured by evaluating the involvement of protein residues (or current flow betweenness) in the allosteric pathways. Present calculations show lower current flow betweennesses of protein residues in presence of lipids as compared to those of the standalone protein. This suggests that the information flow along allosteric communications between EC region and aromatic A-loop residue propagates across the lipid bilayer (Supplementary Fig. 14). Interestingly, efficiencies of distant EC to A-loop pathways are systematically lower in OF *b*MRP1-ATP-ADP than in OF *b*MRP1-(ATP)_2_ (Fig. 6d). Similar trends were observed while considering crossed allosteric communications (Supplementary Fig. 14), *i.e*., EC1 to NBS2 and EC2 to NBS1 pathways. This strongly support the hypothesis that the distant communications between EC and IC regions is lowered after ATP hydrolysis and phosphate release.

## 4. Discussion

*b*MRP1 is the only drug exporter in ABCC family for which there are resolved cryo-EM structures available, so far. The ITC^5,9,60^ has stressed out their clinical and pharmacological relevance. Knowledge about MRP1 dynamics and functions is still fragmented, in contrast to its cousin ABCB1/P-gp. In the present work, all-atom MD simulations were performed in the pre- and post-hydrolysis states, respectively OF *b*MRP1-(ATP)_2_ and OF *b*MRP1-ATP-ADP, in POPC:POPE:Chol (2:1:1) lipid bilayer model to compare their dynamics and structures as well as the putative interplay with surrounding lipids.

Our results strengthen the expected central role of cholesterol involving the so-called CRAC and CARC motifs. These motifs were also observed in other ABC transporters as for MRP1^52^. Our simulations confirmed their importance for cholesterol binding events, even though other sequences might also be involved in line with observations made in the interplay between P-gp and the lipid bilayer^52^. We recently showed that cholesterol molecules bound to pre-TMH1 and pre-TMH7 regions were actively involved in the distant communication between substrate binding pocket and NBSs^14^. Similar trends were observed in the present study in which, annular lipids significantly improve the distant communication between EC and IC regions of *b*MRP1. One can here hypothesize that out of the physical role of a lipid bilayer to maintain the TMH arrangement in the membrane, surrounding lipids might also act as allosteric partners by binding ABC protein to potentiate allosteric communications along and between TMDs. Besides, out of the role of the membrane and despite the small overall conformational difference between pre- and post-hydrolysis states, the latter exhibited a lower efficiency regarding the information flow of EC-to-IC allosteric communications. This might suggest that once the substrate has been released and the nucleotide has been hydrolysed, the IC region may be less sensitive to events occurring on the EC side of the membrane. We also paid particular attention to the putative structural dynamics of the L_0_ domain. MD simulations suggest the existence of a dynamic interaction with NBD1, which tends to be more pronounced in the post-hydrolysis state. This can be correlated with structural observations showing that L_0_ domain interacts with the R-domain connecting NBD1 to TMH7 in the Ycf1p ABCC homolog^20^. Even though no clear clue was obtained to better understand the role of L_0_, previous experiments have shown that the lasso motif is required for MRP1 function^1,6,20^ and its post-translational modifications (*e.g*., phosphorylation) might be involved in MRP1 function regulation, as shown for a human MRP1 ortholog^61^. Our simulations showed that L_0_ flexibility is lowered in the post-hydrolysis state, especially for the L_0_ disordered region, likely owing to the aforementioned interactions with NBD1. Altogether with our present results, this may suggest that the L_0_-NBD1 non-covalent interactions could play a role in triggering the OF-to-IF large scale conformational changes. However, in the present model, no post-translational modification of L_0_ was considered while it may play a role in the L_0_ dynamics. Given that L_0_ may also be dynamically in contact with lipid polar head groups, one can expect that the post-translational phosphorylation of L_0_ might also modulate contacts with inner leaflet which is rich in anionic lipids (*e.g*., phosphoserine, phosphatidic acid polar head). However, further joint experimental and computational structural investigations taking into account the whole system should be considered. This is particularly true since it has recently been suggested that the phosphorylation status of Ycf1p L_0_ strongly depends on the exposure of Ser251 and thus L_0_ dynamics^20^.

Overall, the global conformational dynamics of OF systems in POPC:POPE:Chol (2:1:1) show unfavourable EC opening, smaller NBD twist, and higher NBD distance. Such trends are more pronounced in the post-hydrolysis state indicating that it adopts a closed-cleft (cc) conformation rather than an outward-open conformation. EC closing and IC opening are likely necessary for the OF-to-IF conformational changes. Present results as well as our previous study^14^ clearly indicate that outward-facing open conformation is a transient state which may exist only for the substrate release. We here, therefore, suggest that the use of “OF” conformation for such states should perhaps be replaced by the so-called cc-conformation as proposed for ABCB4^62^. Noteworthy, only a few IV-fold ABC transporters were resolved with a clear open OF conformation (*e.g*., the historical Sav1866^63^ or TmrAB^17^) as shown in Supplementary Fig. 15. Considering the ABC conformational space, two populations for OF *b*MRP1-ATP-ADP state were observed, for which transition would require IC closing. Remarkably, the conformation exhibiting the smallest EC angle was observed only in one replica, even though all replicas started from the same initial structure at the pre-equilibration step. Therefore, the existence of such conformation should be considered very carefully.

NBD rocker-switch angle slightly but significantly deviates by *ca*. 5° between pre- and post-hydrolysis states. In agreement with structural insights observed in P-gp, hydrolysis induces local conformational changes in the conserved motifs, mostly in the signature motif^64^. Besides, we here propose that ATP hydrolysis also affects the local Walker B conformation which participates in Mg^2+^-binding. However, the present model considered anionic catalytic glutamate instead of glutamic acid protonation state. Given the chemical and water environment, one can expect that the deprotonation event might occur rapidly after ATP-hydrolysis. In agreement with studies carried out on P-gp in which EC opening event was correlated to distant A-loop position^11^, our results confirmed the long distance communication between IC and EC regions. However, in *b*MRP1, the calculated EC opening deviation remains lower than in P-gp. This might be explained by a different inherent behaviour of MRP1, which may be more pronounced owing to the degenerate NBD1. Indeed, as shown in our former study^14^, NBD1 coupling helix does not adopt the “ball-and-socket” conformation, leading to a lower signal transduction from NBD1 to TMDs. We here confirm that ATP-binding in NBS1 likely acts as a pivot since NBD1 and NBS1 are almost structurally identical regardless of the hydrolysis state. NBD2 exhibit pseudo-asymmetric structural deformation on the NBS2 side after ATP-hydrolysis. This was confirmed by monitoring local distances, as well as non-covalent interactions between magnesium atoms, nucleotides, and protein residues (including H-bond, van der Waals and Coulomb potentials). We are in agreement with the description stronger interactions between the ATP and conserved motifs in NBS1 than in NBS2 even in the post-hydrolysis state. This strengthens the hypothesis suggesting that ATP release from NBS1 is less likely^15^.

Overall, the OF *b*MRP1-ATP-ADP state is not expected to be a resting state since it might undergo ADP phosphate release. We here discussed about OF *b*MRP1-ATP-ADP state and not UR conformation since (i) projection of *b*MRP1 IC angles onto the ABC conformational space was distant to those calculated from cryo-EM asymmetric TmrAB UR conformations^17^ and (ii) overall conformations of *b*MRP1 cryo-EM structures (6UY0 and 6BHU) are very similar^12^. Given that no NBD dissociation was observed, our MD simulations are in line with the hypothesis that the rate limiting step of OF-to-IF transition is the nucleotide (ADP) release event which might be associated with NBD dissociation^12^. However, the absence of significant differences regarding ATP-binding mode in NBS1 of post-hydrolysis state as compared to pre-hydrolysis support that only a partial NBD dissociation may occur. This might be associated with the ATP molecule that remains bound along multiple transport cycles, as proposed for NBS degenerate ABC transporters^15^.

## Supporting information

Electronic Supplementary information

## 5. Acknowledgements

We thank Xavier Montagutelli from the IT department of Limoges University for technically supporting supercomputer facilities. Calculations were performed using CALI (“CAlcul en LImousin”) and “Baba Yaga” supercomputers hosting in Limoges University, as well as on the “Jean-Zay” national supercomputer from IDRIS HPC resources under the allocations 2020-A0080711487, 2021-A0100711487 and 2022-A0120711487 made by GENCI. This work was supported by grants from the “Agence Nationale de la Recherche” (ANR-19-CE17-0020-01 IMOTEP and ANR-21-CE18-0030 RAPRACLID), Région Nouvelle Aquitaine and “Institut National de la Santé et de la Recherche Médicale” (INSERM, AAP-NA-2019-VICTOR).

## 7. Authorship

Á.T and F.D.M conceived the study. Á.T and F.D.M. conducted all the simulations. Á.T, A.J., V.C and F.D.M analysed the simulations. Á.T. and F.D.M. interpretated the results and discussed them together with A.J. and V.C. Á.T. and F.D.M wrote the initial draft, and it was edited, reviewed, and approved by all authors.

## 8. Data availability

All data analysed during this study are included in this published article (and its supplementary information files. MD trajectories are available upon reasonable requests.

