## Supplementary material for "Computational and structural insights into the pre- and post-hydrolysis states of bovine multidrug resistance associated protein 1 (MRP1)": Electronic Supplementary information

*Inserm UMR 1248 Pharmacology & Transplantation, Univ. Limoges, 2 rue du Prof. Descottes, 87000 F-Limoges, France*

**Contact information**
Dr. Florent Di Meo & Dr. Ágota Tóth
Inserm U1248 Pharmacology & Transplantation, Univ. Limoges,
2 rue du Prof. Descottes,
87000 F-Limoges, France


### List of Supplementary Figures

**Supplementary Figure 1. Root-mean-square deviation (RMSD) along MD simulations calculated for the pre- and post-hydrolysis states.** Replica 1, 2 and 3 are coloured red, blue and yellow, respectively. 7

**Supplementary Figure 2. Root-mean-square deviation (RMSD) excluding the lasso domain ($L_0$) along MD simulations calculated for the pre- and post-hydrolysis states.** Replica 1, 2 and 3 are coloured red, blue and yellow, respectively. 7

**Supplementary Figure 3. IC angle (°) during the whole simulations calculated for the pre- and post-hydrolysis states.** Replica 1, 2 and 3 are coloured red, blue and yellow, respectively. 8

**Supplementary Figure 4. Evolution of EC angle (°) along MD simulations calculated for the pre- and post-hydrolysis states.** Replica 1, 2 and 3 are coloured red, blue and yellow, respectively. 8

**Supplementary Figure 5. EC distance (Å) during the whole simulations calculated for the pre- and post-hydrolysis states**. Replica 1, 2 and 3 are coloured red, blue and yellow, respectively. 9

**Supplementary Figure 6. NBD distance (Å) during the whole simulations calculated for the pre- and post-hydrolysis states.** Replica 1, 2 and 3 are coloured red, blue and yellow, respectively. 9

**Supplementary Figure 7. NBD twist angle (°) during the whole simulations calculated for the pre- and post-hydrolysis states.** Replica 1, 2 and 3 are coloured red, blue and yellow, respectively. 10

**Supplementary Figure 8. Convergence of InfleCS calculations for the post-hydrolysis sates.** 10

**Supplementary Figure 9. Contacts between the nucleotides and their binding site.** Conserved motives are shown by different colours: Walker A red, Walker B blue, signature motif green, A-loop orange, Q-loop teal, X-loop pink, D-loop cyan, and H-loop yellow. 11

**Supplementary Figure 10. PE-lipid distribution of pre- and post-hydrolysis states.** 12

**Supplementary Figure 11. Cholesterol distribution of pre- and post-hydrolysis states.** 13

**Supplementary Figure 12. Non-covalent interaction energies between nucleotides and NBSs extracted from Coulomb (top) and van der Waals (bottom) potentials obtained from the performed MD simulations.** 14

**Supplementary Figure 13. Calculated per-residue betweenness** in the allosteric pathway from extracellular helices to NBS1 and NBS2 A-loop aromatic residue for **a)** pre-hydrolysis and **b)** post-hydrolysis states. 15

**Supplementary Figure 14. Allosteric efficiency from the EC regions to crossed NBS A-loop aromatic residues, namely EC1 to A-loop NBS2 (left) and EC2 to A-loop NBS1 (right).** 16

**Supplementary Figure 15. Sav1866 (PDBID: 2HYD) in OF open conformation.** 17

### Supplementary Tables

**Supplementary Table 1. $L_0$ modelling of OF states.**

| | |
|---|---|
| 269-RKQPVKIV-276 | $L_0$ of IF system |
| 277-YSSKDPAKPKGSSKVDV-293 | sequence |
| 294-NEEAEALIVKCPQKERD-310 | $L_0$ of IF system |

**Supplementary Table 2. The box size (Å$^3$) of pre- and post-hydrolysis states.**

| | **POPC:POPE:Chol (2:1:1)** |
|---|---|
| pre-hydrolysis | 123.1x126.3x179.5 |
| post-hydrolysis | 122.0x121.8x180.1 |

**Supplementary Table 3. The number of atoms investigated in the present study.**

| | **POPC:POPE:Chol (2:1:1)** |
|---|---|
| pre-hydrolysis | 245 145 |
| post-hydrolysis | 241 635 |

**Supplementary Table 4. ABC structural parameters** (EC angle, IC angle, NBD distance and NBD twist, values for a set of selected, resolved ABC type IV proteins).

| PDB ID | EC angle (°) | IC angle (°) | NBD distance (Å) | NBD twist (°) |
|---|---|---|---|---|
| 5UJ9 | 15.74 | 36.75 | 55.41 | -158.29 |
| 5UJA | 15.27 | 29.59 | 43.3 | -149.74 |
| 6BHU | 16.7 | 21.57 | 27.47 | -149.61 |
| 6UY0 | 17.21 | 21.27 | 28.03 | -149.6 |
| 4PL0 | 12.97 | 22.16 | 27.57 | -146.93 |
| 5TTP | 13.12 | 21.24 | 28.26 | -159.91 |
| 4RY2 | 15.06 | 27.84 | 32.98 | -143.78 |
| 4S0F | 15.5 | 20.84 | 27.61 | -151.14 |
| 5C73 | 19.53 | 21.93 | 27.96 | -152.1 |
| 2ONJ | 21.08 | 21.65 | 27.43 | -155.02 |
| 4Q4A | 14.73 | 35.49 | 34.49 | -147.15 |
| 5MKK | 15.16 | 33.12 | 38.93 | -139.16 |
| 6RAF | 14.97 | 29.88 | 39.01 | -139.81 |
| 6RAG | 15.35 | 32.56 | 39.83 | -140.99 |
| 6RAH | 22.4 | 21.19 | 27.92 | -155.04 |
| 6RAI | 13.59 | 22.3 | 27.97 | -156.89 |
| 6RAJ | 24.03 | 21.68 | 28.02 | -156.08 |
| 6RAK | 14.12 | 22.07 | 28.01 | -157.07 |
| 6RAL | 14.48 | 23.31 | 28.77 | -155.81 |
| 6RAM | 14.7 | 25.37 | 29.45 | -156.16 |
| 6RAN | 14.78 | 33.55 | 40.46 | -143.04 |
| 6LR0 | 14.83 | 41.34 | 54.72 | -147.24 |
| 6S7P | 15.67 | 20.99 | 28.1 | -153.66 |
| 6C0V | 15.91 | 20.86 | 28.77 | -157.25 |
| 6QEX | 12.87 | 34.16 | 38.41 | -144.65 |

**Supplementary Table 5. Selected residues for allosteric pathway calculations from extra-cellular helices to NBS1 and NBS2.**

| **Sink A-loop aromatic residue** | | **Source** | |
|---|---|---|---|
| **NBS1** | **NBS2** | **TMH1, 2, 10 and 11** | **TMH4, 5, 7 and 8** |
| Trp653 | Tyr1301 | Leu348<br>Leu349<br>Ile350<br>Asn351<br>Phe352<br>Glu360<br>Trp361<br>Gln362<br>Gly363<br>Arg1221<br>Val1229<br>Gly1230<br>Leu1231<br>Ser1232<br>Val1233 | Leu460<br>Phe565<br>Ala566<br>Val567<br>Tyr568<br>Val569<br>Lys581<br>Ala582<br>Phe583<br>Val584<br>Ser585<br>Trp998<br>Thr999<br>Asp1000<br>Asp1001<br>Arg1014<br>Leu1015<br>Ser1016<br>Val1017<br>Tyr1018<br>Gly1019 |

### Supplementary Figures

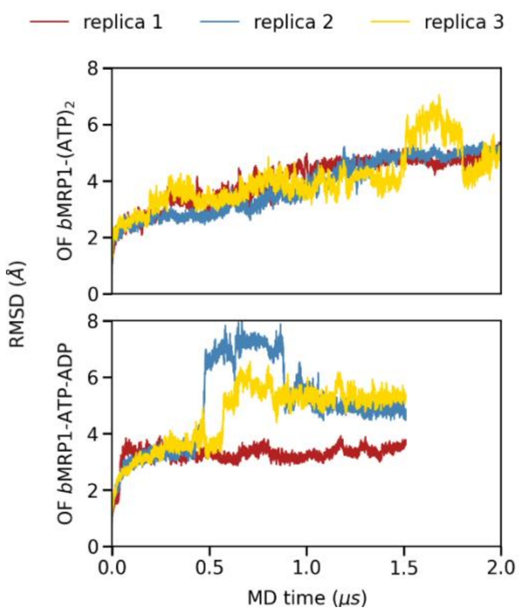

**Supplementary Figure 1. Root-mean-square deviation (RMSD) along MD simulations calculated for the pre- and post-hydrolysis states.** Replica 1, 2 and 3 are coloured red, blue and yellow, respectively.

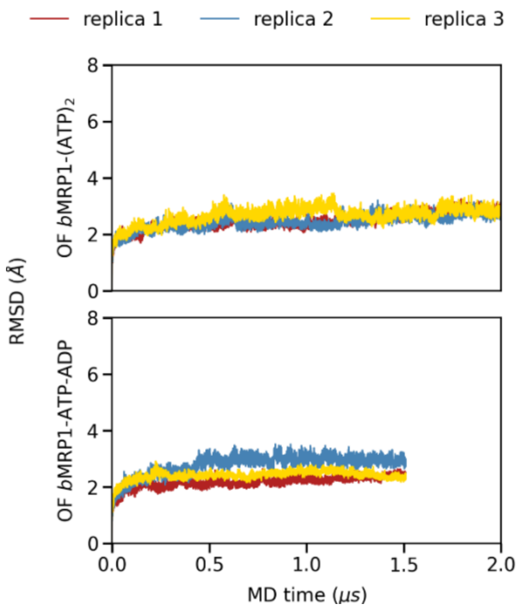

**Supplementary Figure 2. Root-mean-square deviation (RMSD) excluding the lasso domain ($L_0$) along MD simulations calculated for the pre- and post-hydrolysis states.** Replica 1, 2 and 3 are coloured red, blue and yellow, respectively.

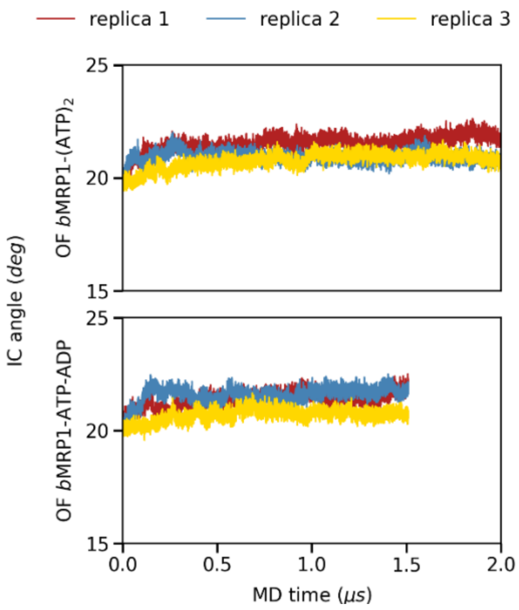

**Supplementary Figure 3. IC angle (°) during the whole simulations calculated for the pre- and post-hydrolysis states.** Replica 1, 2 and 3 are coloured red, blue and yellow, respectively.

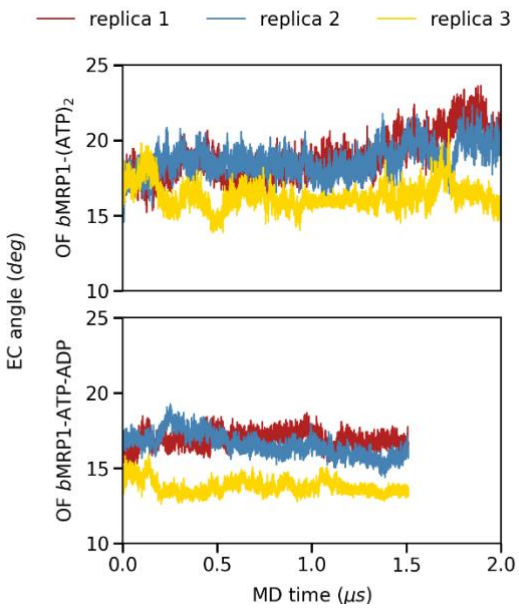

**Supplementary Figure 4. Evolution of EC angle (°) along MD simulations calculated for the pre- and post-hydrolysis states.** Replica 1, 2 and 3 are coloured red, blue and yellow, respectively.

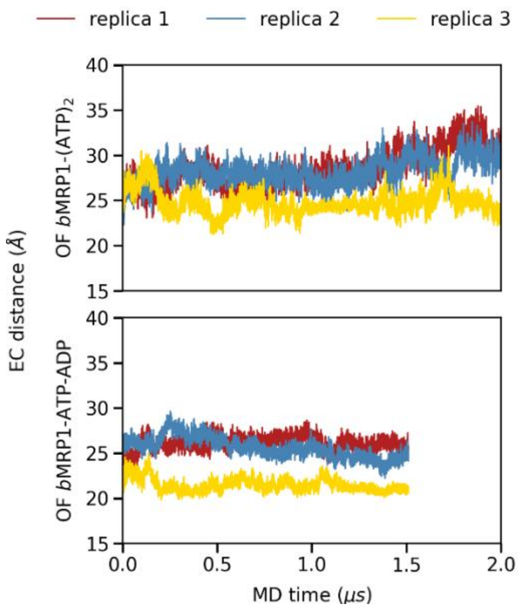

**Supplementary Figure 5. EC distance (Å) during the whole simulations calculated for the pre- and post-hydrolysis states**. Replica 1, 2 and 3 are coloured red, blue and yellow, respectively.

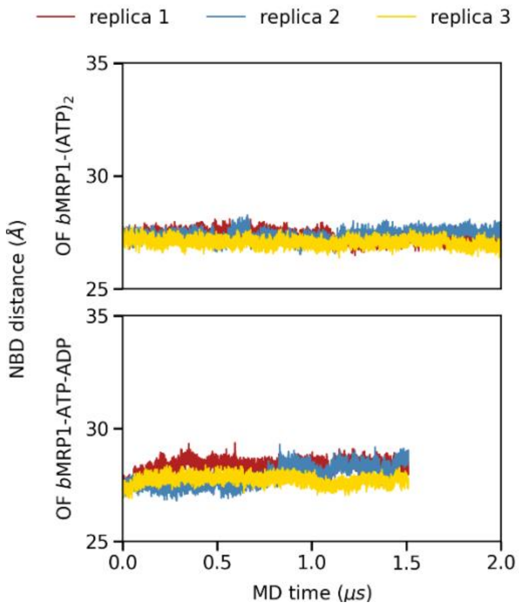

**Supplementary Figure 6. NBD distance (Å) during the whole simulations calculated for the pre- and post-hydrolysis states.** Replica 1, 2 and 3 are coloured red, blue and yellow, respectively.

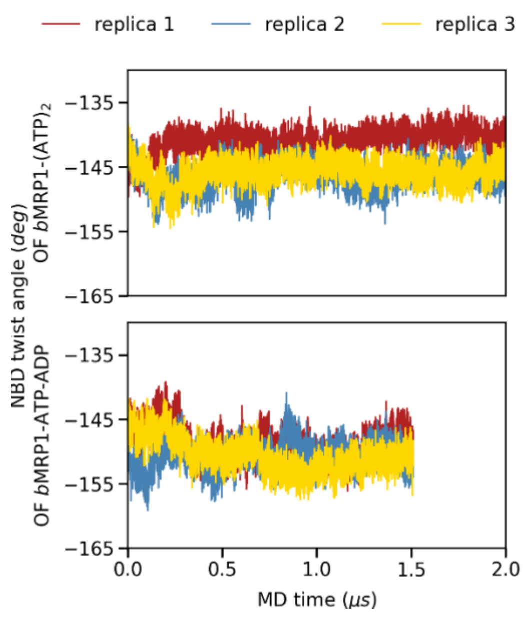

**Supplementary Figure 7. NBD twist angle (°) during the whole simulations calculated for the pre- and post-hydrolysis states.** Replica 1, 2 and 3 are coloured red, blue and yellow, respectively.

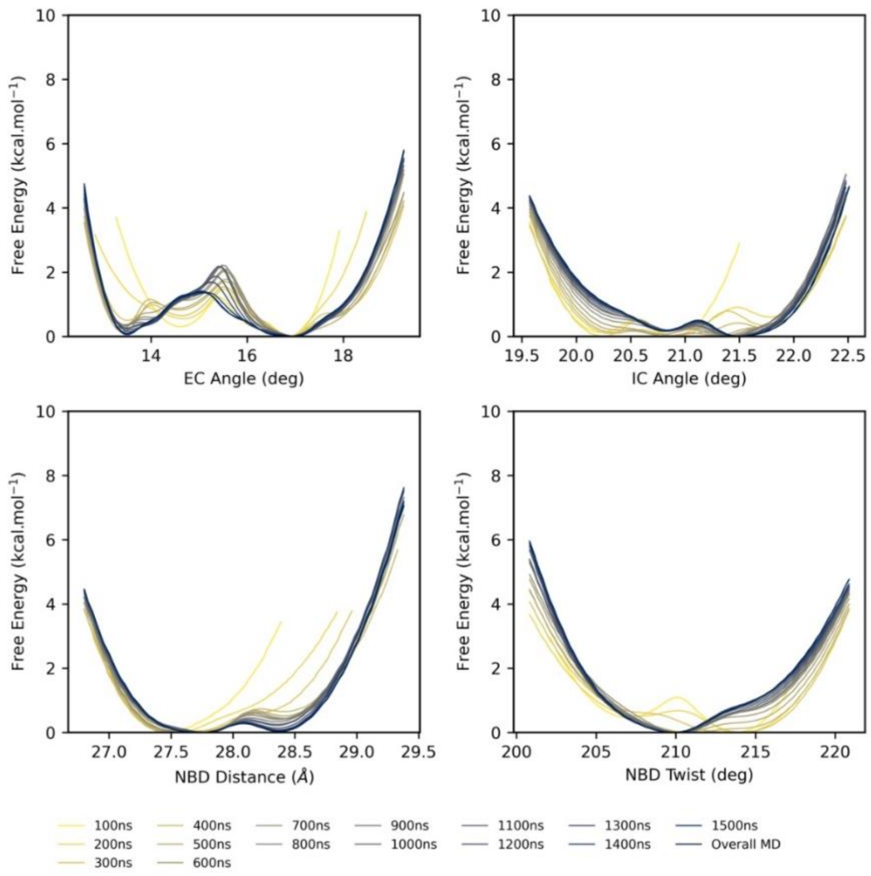

**Supplementary Figure 8. Convergence of InfleCS calculations for the post-hydrolysis sates.**

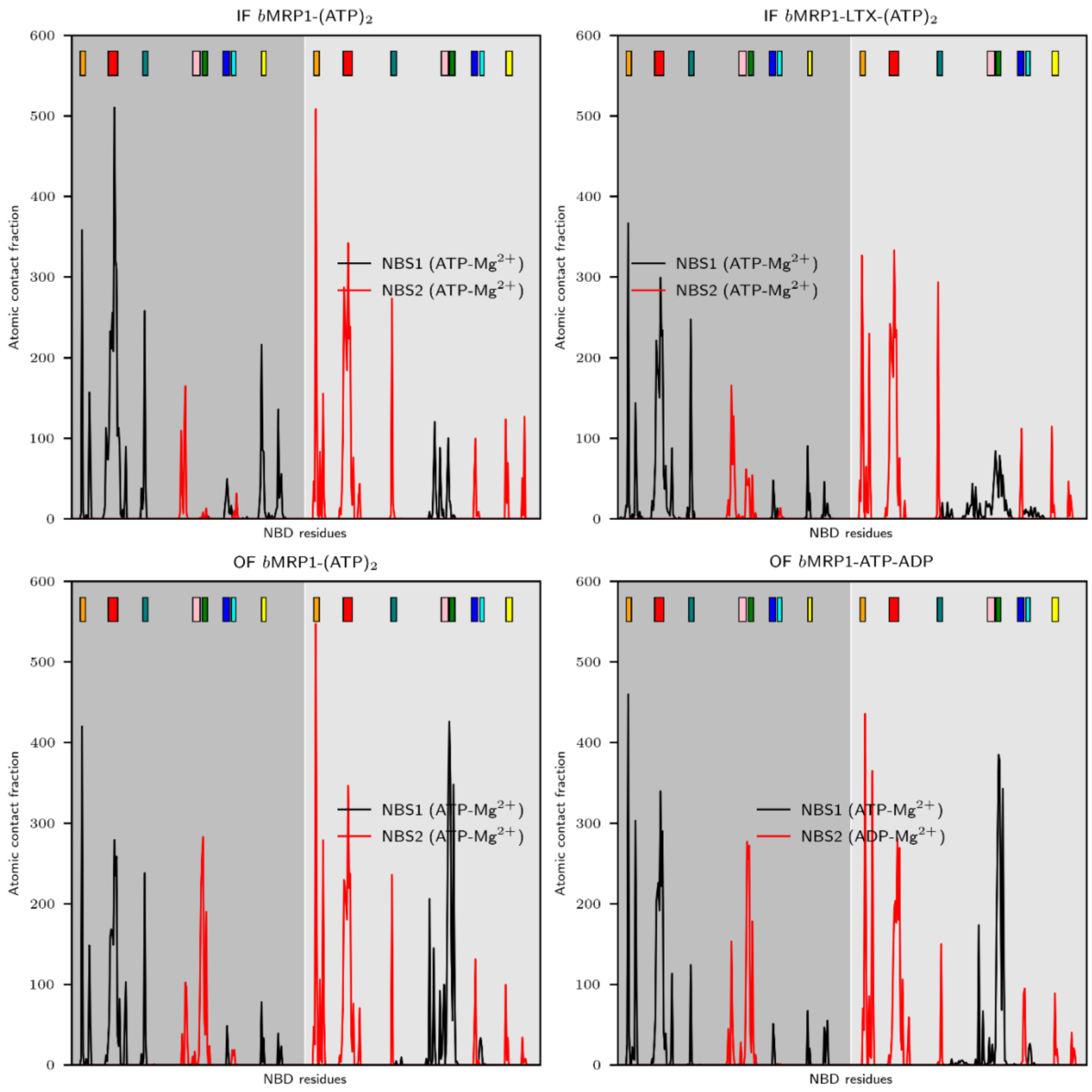

**Supplementary Figure 9. Contacts between the nucleotides and their binding site.** Conserved motives are shown by different colours: Walker A red, Walker B blue, signature motif green, A-loop orange, Q-loop teal, X-loop pink, D-loop cyan, and H-loop yellow.

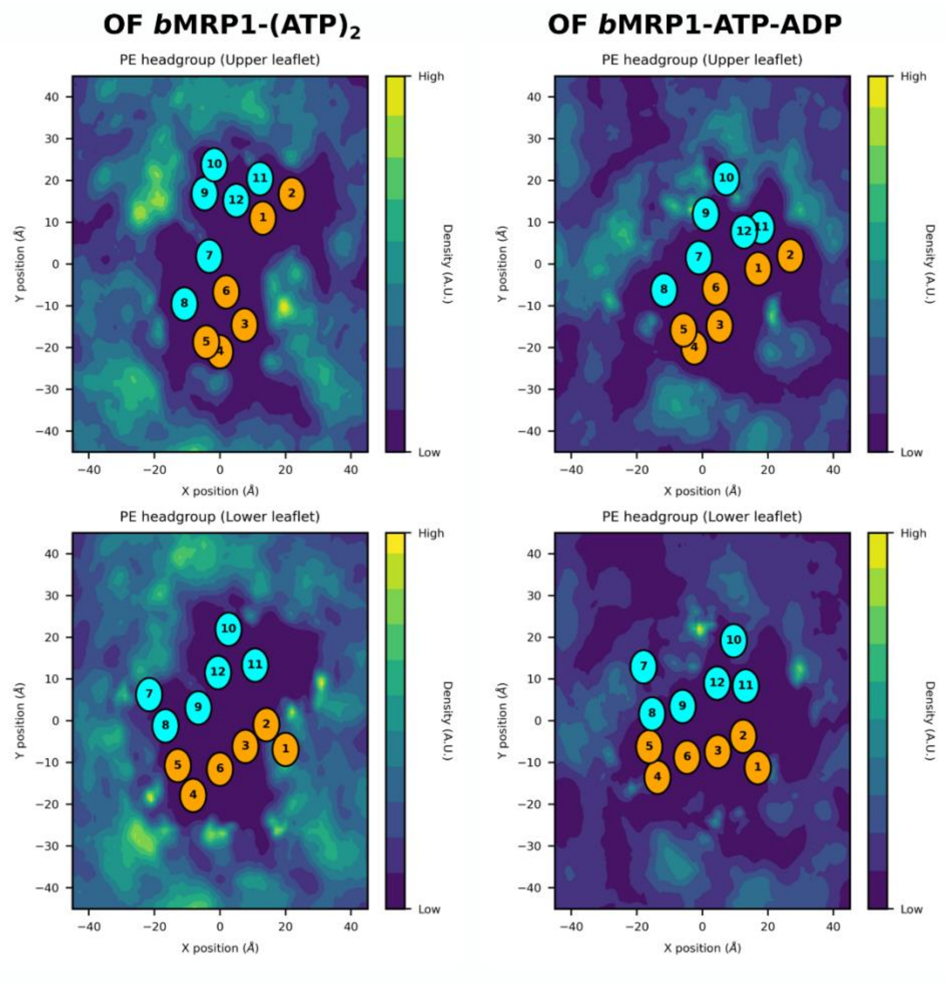

**Supplementary Figure 10. PE-lipid distribution of pre- and post-hydrolysis states.**

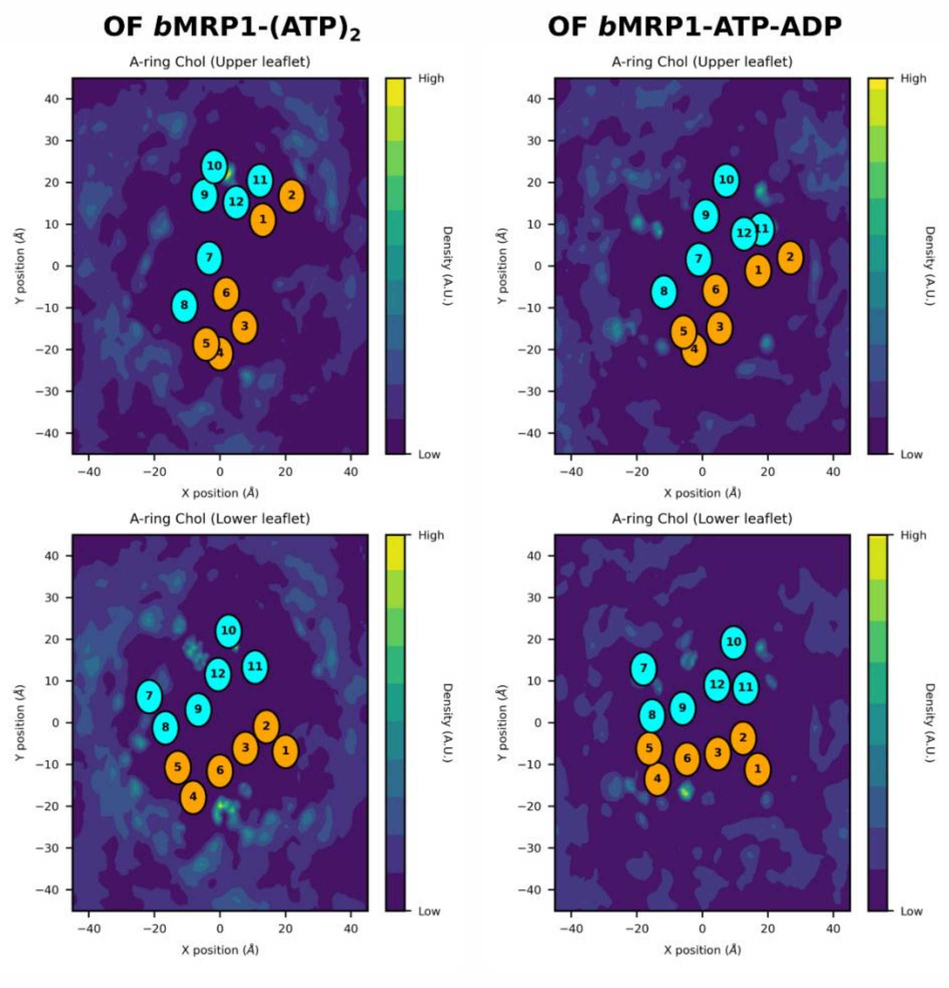

**Supplementary Figure 11. Cholesterol distribution of pre- and post-hydrolysis states.**

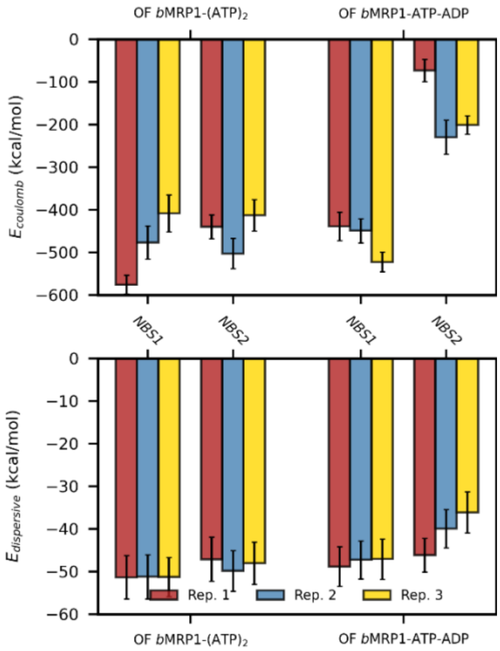

**Supplementary Figure 12. Non-covalent interaction energies between nucleotides and NBSs extracted from Coulomb (top) and van der Waals (bottom) potentials obtained from the performed MD simulations.**

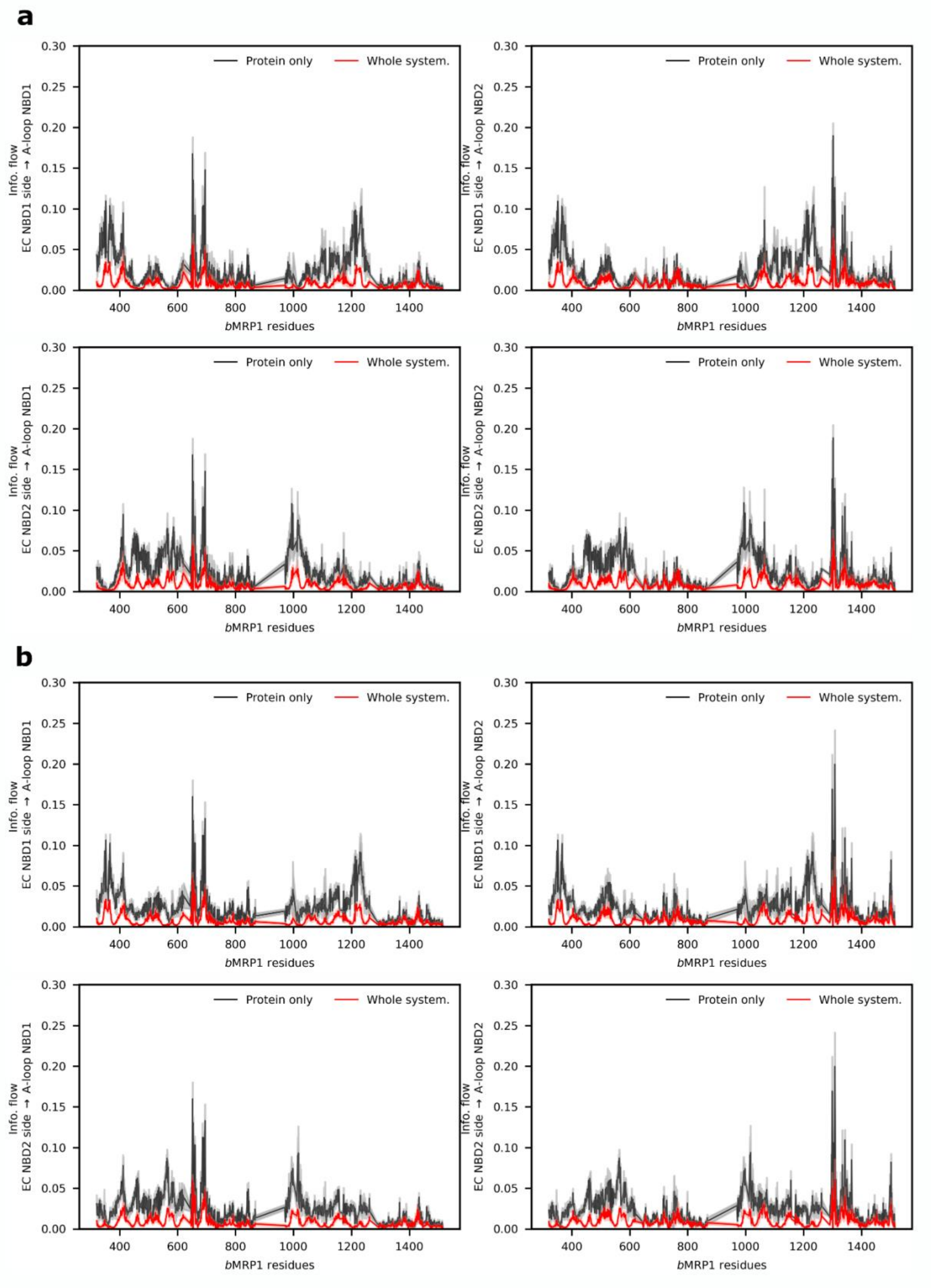

**Supplementary Figure 13. Calculated per-residue betweenness** in the allosteric pathway from extracellular helices to NBS1 and NBS2 A-loop aromatic residue for **a)** pre-hydrolysis and **b)** post-hydrolysis states.

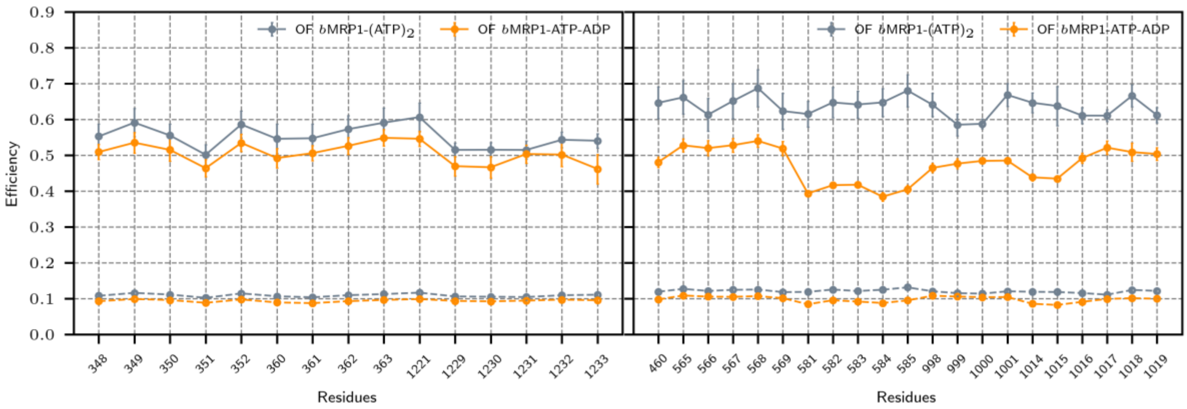

**Supplementary Figure 14. Allosteric efficiency from the EC regions to crossed NBS A-loop aromatic residues, namely EC1 to A-loop NBS2 (left) and EC2 to A-loop NBS1 (right).**

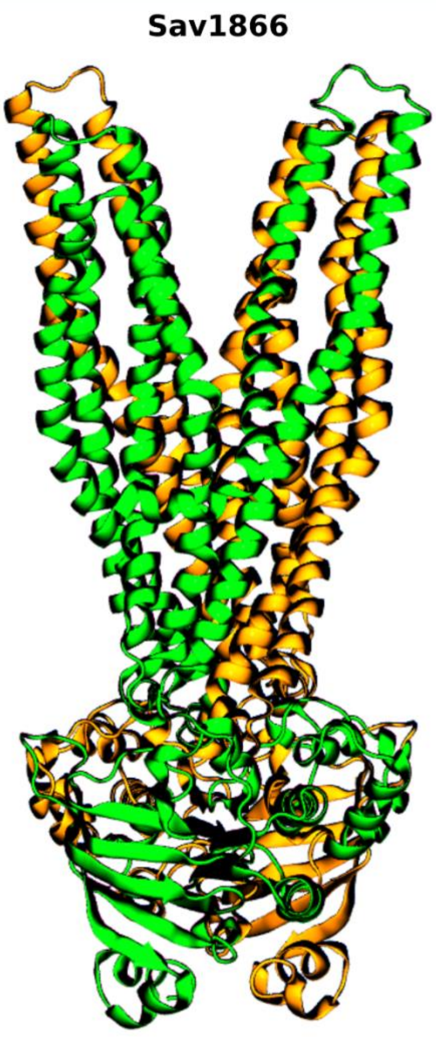

**Supplementary Figure 15. Sav1866 (PDBID: 2HYD) in OF open conformation.**
